# Scalable Condition-relevant Cell Niche Analysis of Spatial Omics Data with Taichi

**DOI:** 10.1101/2024.05.30.596656

**Authors:** Yan Cui, Zhiyuan Yuan

## Abstract

Tissues are composed of heterogeneous cell niches, which can be investigated using spatial omics technologies. Large consortia have accumulated vast amounts of spatially resolved data, which typically assign slice-level condition labels without considering intra-slice heterogeneity, particularly differential cell niches that respond to certain perturbations. Here, we present Taichi, an efficient and scalable method for condition-relevant cell niche analysis that does not rely on pre-defined discrete spatial clustering. Taichi utilizes a scalable spatial co-embedding approach that effectively accounts for batch effects, incorporating advanced label refinement and graph heat diffusion techniques to explore condition-relevant cell niches across extensive multi-slice and multi-condition spatial omics datasets. Comprehensive benchmarks demonstrate Taichi’s ability to precisely identify condition-relevant niches under various levels of perturbations. We showcase Taichi’s effectiveness in accurately delineating major shifts in cell niches in a mouse model of diabetic kidney disease compared to a normal group, revealing disease-specific cell-cell interactions and spatial gene expression patterns. Furthermore, Taichi can identify key subtype-relevant niches between colorectal cancer patient groups with significantly different survival outcomes. Moreover, we demonstrate that Taichi can help discover more fine-grained clinical properties within the originally coarse-defined patient groups in large-scale tumor spatial atlases, reflecting intra-group heterogeneity obscured previously. Additionally, we combine Taichi and tensor decomposition to discover higher-order biomarkers relevant to the immunotherapy response of triple-negative breast cancer. Finally, we highlight Taichi’s speed and scalability by confirming its unique applicability in large-scale scenarios containing up to 16 million cells in ∼ 12 minutes. Taichi provides a powerful tool for mining disease-relevant spatially resolved insights in the era of big data in spatial biology.

## Introduction

Spatial omics technologies have significantly advanced our ability to investigate tissue spatial biology by allowing the simultaneous measurement of molecular and spatial information within tissues^1^. These technologies generate highly multiplexed spatial maps of transcriptomics^2,3^, proteins^4,5^, and/or metabolic features^6,7^ at up to single-cell resolution^8^. Spatial omics technologies surpass traditional single-cell omics by maintaining the spatial context, which is essential for unraveling the complexities of biological processes such as development and disease^9–12^.

In high-throughput omics studies, the comparison of datasets collected under various conditions is crucial for quantifying the impacts of different perturbations and conditions, such as disease, drug treatments, or gene modifications^13^. In single-cell omics, two primary classes of methods are employed to assess the effects of perturbations to identify condition-relevant cell types. The first class relies on pre-defined cell types to quantify shifts in cell type abundance across conditions^14–16^. The second class involves differential abundance analysis^17–20^, which estimates the relative likelihood of cells within cell state space, quantifies relative abundance in transcriptomically defined cellular neighborhoods, or employs other related strategies. However, in current tissue studies, the focus often shifts to multi-cellular organizations or cell niches^21–23^, which are spatially organized regions that respond collectively to perturbations rather than on an individual cell basis^24^. Methods designed for single-cell data typically overlook spatial information, limiting the exploration of higher-level cell niche differences across conditions.

In disease study, accurately identifying and analyzing those perturbed cell niches (i.e., disease-relevant cell niches) is critical. However, typical spatial omics studies often analyze entire tissue slices and categorize them based solely on the meta-information of the patient (e.g., patient’s overall diagnosis), ignoring the variability within the slices themselves^25–27^. For example, early-stage disease like liver fibrosis may exhibit regions that resemble healthy tissue alongside regions showing disease characteristics, but the common practice is to label the entire slice as a “liver fibrosis slice”, resulting in a noisy label problem^6^. Analyzing such noisy-labeled slices without differentiating between healthy and disease-relevant areas can significantly weaken the power of the statistical analysis used to understand disease. This issue becomes even more complex in diseases like cancer, where tumors might exhibit a mix of characteristics shared across or specific to different subtypes or a complex hierarchy of niches that affect disease progression and treatment responses^27,28^.

In spatial omics community, computational methods that consider both spatial and molecular profiles generally concentrate on spatial clustering to stratify cell niches within individual slice^29–31^, with a few exceptions that are capable of identifying cell niches across multiple slices^32–34^, leaving the identification of condition-relevant cell niches largely unaddressed. The few existing cross-slice methods that can identify niches with significant differences in abundance often rely on pre-defined discrete spatial clusters, which may not accurately capture the continuous nature of spatial contexts within tissues and may introduce biases based on prior knowledge (e.g., the number of spatial clusters in the tissue). These methods also typically require multiple biological replicates to determine the statistical significance of niche abundance differences, which can be impractical when dealing with rare or limited samples^35,36^. Furthermore, as spatial omics data become increasingly available through various consortia efforts, batch effects across slices and the scalability limitations of current spatial clustering methods pose significant challenges for analyzing large-scale heterogeneous spatial datasets^27,37,38^.

To address these challenges, we developed Taichi, an efficient and scalable method for condition-relevant cell niche (CRN) analysis that does not rely on pre-defined spatial clustering for discrete spatial domains. Taichi employs a scalable spatial co-embedding approach, coupled with label refinement and graph heat diffusion techniques, to effectively explore condition-relevant cell niches across extensive multi-slice and multi-condition spatial omics datasets. By comparing healthy and disease conditions, Taichi partitions each disease slice (potentially containing a mixture of healthy and disease regions) into two virtual slices representing health and disease, facilitating the discovery of spatially resolved insights in purer disease-relevant niches. Furthermore, when comparing different disease subtypes, Taichi identifies the key cell niches that define disease subtypes, potentially uncovering novel insights through in-depth analysis.

To the best of our knowledge, Taichi is the first method capable of identifying condition-relevant cell niches at scale without pre-defined spatial clustering. We benchmarked Taichi against other methods using meticulously designed simulation datasets to identify different levels of spatial perturbations. Our results demonstrate Taichi’s effectiveness in accurately delineating major shifts in cell niches in a mouse model of diabetic kidney disease (DKD). With the shift in cell niches, we further reveal changes in cell type proportions and DKD-related cell-cell interaction patterns. Additionally, we identify key DKD-relevant niche-associated spatial domain-specific genes and metagenes, demonstrating the reliability of Taichi outcomes in resolving gene expression mechanisms. Moreover, we distinguish key group-specific niches between patient groups in colorectal cancer (CRC), resulting in significantly different survival outcomes and more meaningful differential cellular neighborhoods than the original coarse labels. Taichi also helps discover more fine-grained and multi-aspect clinical properties (e.g., different recurrence, subtype situation) within original coarse-defined patient group labels in large-scale tumor spatial atlases. Furthermore, we show that Taichi can discover high-order biomarkers related to immunotherapy response of triple-negative breast cancer (TNBC) with tensor decomposition. Finally, we highlight Taichi’s speed and scalability, confirming its unique applicability in two forms of extremely large-scale spatial omics datasets using limited time and computational resources.

## Results

### The noisy labeling problem

Taichi is introduced on the principle that each tissue slice comprises a collection of cell niches, and not all niches within a slice are necessarily related to the slice’s condition label. For instance, a slice from a diseased individual may contain both healthy and disease-relevant niches. Consequently, we consider the traditional approach of assigning condition labels to each slice (e.g., diseased or normal) as a form of noisy labeling problem. The primary objective of Taichi is to refine these noisy labels to identify true differences in cell niches across conditions.

The input of Taichi is spatial omics data with slice-level condition labels (Fig. 1a (i)). The initial step in the Taichi workflow is to construct a shared niche embedding space across conditions (e.g., different disease status, development stages, etc.). This shared embedding space is expected to represent each niche by simultaneously capturing information about the cell state/type and spatial context. To achieve this, Taichi employs a scalable spatial embedding approach, Multi-range cEll coNtext DEciphereR (MENDER)^39^ that harmonize the diverse slices into a batch-free embedding space, capturing both intra-slice heterogeneity and inter-slice variances (Fig. 1a (ii)). In this embedding space, niche manifold may be discrete or continuous. Instead of performing clustering on this embedding space to obtain discrete niche clusters and performing niche cluster-level differential testing between conditions, which may lead to the loss of continuity and require additional prior knowledge of tissue structures, Taichi proceeds to calculate the probability of each cell niche belonging to the condition group based on its spatial embedding and the original, potentially noisy label (Fig. 1a (iii)). By performing graph heat diffusion on the original physical space, Taichi can effectively segregate the tissue slices into refined and continuous spatial niches, indicating differences and consistent niches across conditions (Fig. 1a (iv)). Finally, a simple k-means based segmentation is performed on the diffused result to determine the exact niche-wise condition label (Fig. 1a (v)). This approach allows for the precise delineation of niches significantly impacted by conditions. Taichi has many applications, including comparing healthy and disease samples (Fig. 1a (vi)), comparing disease subtypes (Fig. 1a (vii)), and defining spatial indicators for fine-grained patient stratification (Fig. 1a (viii)). We used 15 datasets with different characteristics to benchmark Taichi’s accuracy, continuity and robustness, showcase Taichi’s biomedical applications, and demonstrate Taichi’s unique scalability on extremely large spatial datasets (Fig. 1b). Details of Taichi are explained in Methods.

**Fig. 1.**
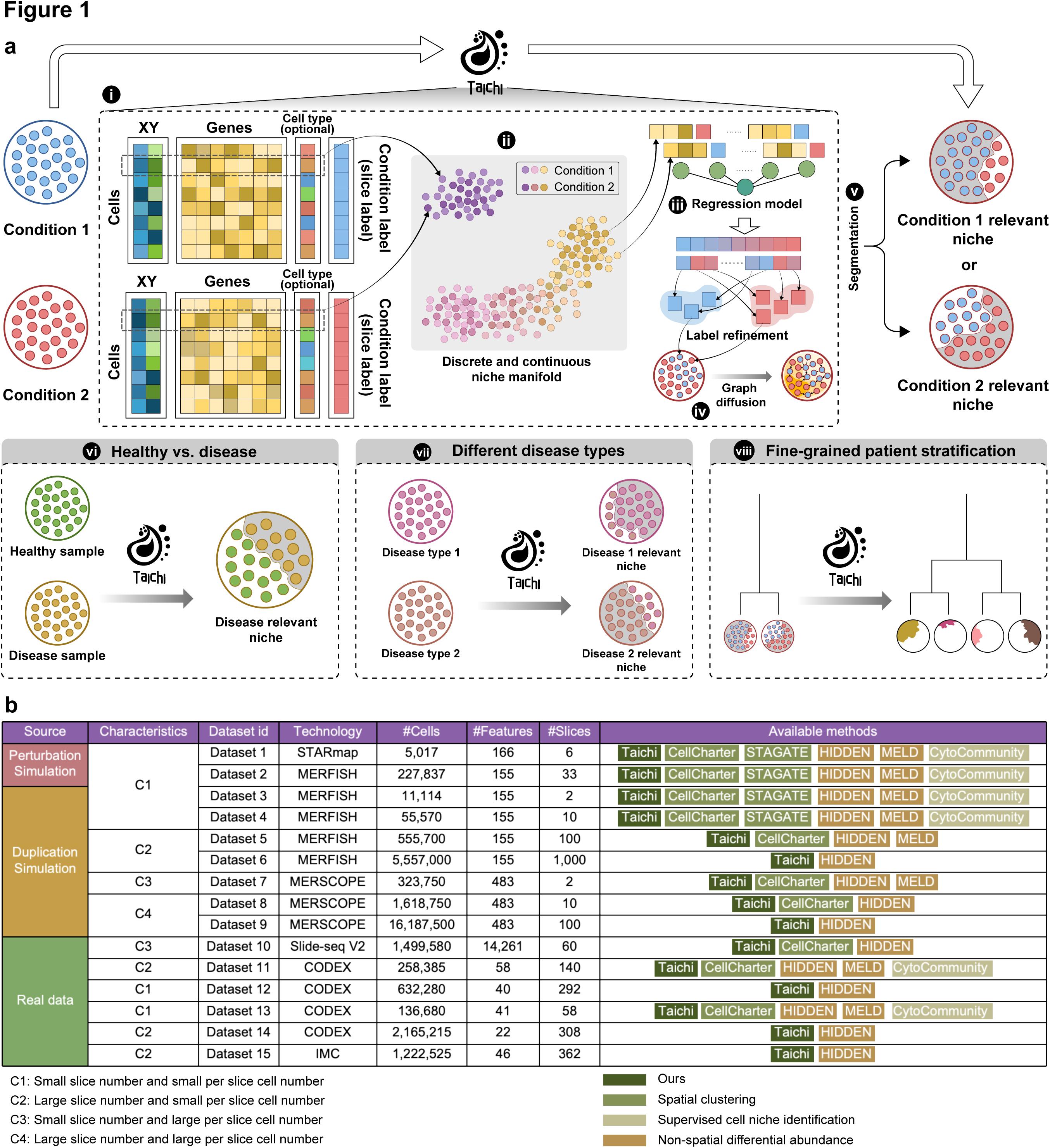
| Workflow of Taichi and datasets. **a,** (i) Workflow of the Taichi process for identifying the condition-relevant niches with slice-level-annotated multi-slice spatial omics data, where each slice is annotated with a slice-level label, e.g., condition and control. The input spatial omics data contain each cell’s expression vector, spatial location, and predefined cell type label (optional). Taichi assigns the cell-level input label based on the slice annotation to which it belongs, which will be refined in the subsequent process. (ii) Cell/spot is embedded into a co-embedding space without batch effects among slices. Specifically, each cell/spot is embedded based on the spatial arrangement of local neighborhood. These embeddings are then utilized in the label refinement process. (iii) With the co-embedding representation vector for each cell, which includes niche information, Taichi applies label refinement to both conditions’ cells/spots. (iv) Taichi transfers the refined label back to corresponding condition slice groups and applies graph heat diffusion based on the spatial neighborhood graph to obtain a spatial-aware, and continuous condition-relevant score. (v) Taichi segment the area in the slice spatial space based on the smoothed scores. (vi) Taichi can be utilized to analyze health and disease information to identify disease-relevant niches. (vii) Taichi can also be used to identify disease subtype-relevant niches within disease groups. (viii) The distribution of the condition-relevant niches can be used for fine-grained annotation within original condition groups. **b.** In our study, we used the adapted state-of-the-art methods from related domains as competing methods, and categorized them into four groups: Spatial clustering methods, non-spatial differential abundance methods, supervised niche identification methods, and ours (Taichi). We partitioned the datasets based on their properties, C1, C2, C3, and C4, covering various scenarios (see “data availability” in Methods). An available method is defined as one that does not encounter out-of-memory errors and has a running time of less than 8 hours on the dataset.

### Benchmarking Design

To evaluate the performance of Taichi in identifying condition-relevant cell niches (CRN) compared to other methods, we conducted a benchmarking analysis using a set of simulation datasets derived from two widely used spatial omics technologies, STARmap^40^ and MERFISH^41^, consisting of 11 condition pairs. Although no existing methods are specifically designed for CRN analysis other than Taichi, we compared two sets of existing methods (see Methods): differential abundance (DA) analysis methods for single-cell (non-spatial) data, including MELD^17^ and HIDDEN^42^, and spatial clustering methods for spatial data, including STAGATE^34^, CellCharter^22^, and CytoCommunity^43^. DA analysis methods are designed for identifying differences in cell state between conditions but do not consider the spatial context of cells. MELD was selected for the recommendation in a recent benchmark study^44^. HIDDEN was selected as one of the most recent DA methods^42^. Although spatial clustering methods are not designed to identify differences between conditions, we implemented additional modules to enable CRN analysis for them (see Methods). STAGATE was selected based on the recommendation in a recent benchmark study^45^. CellCharter^22^ and CytoCommunity^43^ were selected as they represent the most advanced spatial clustering methods developed after 2024. This benchmarking design aims to provide a comprehensive evaluation of Taichi’s performance in identifying CRNs compared to adapted state-of-the-art methods from related domains.

### Identifying Addition Perturbations

We first designed a straightforward simulation to test whether different methods can accurately identify simple perturbations in cell niches, specifically the addition of new cell niches. We simulated the first three condition pairs from real STARmap datasets of the mouse medial prefrontal cortex^40^ (Fig. 2a, see Methods). The real datasets contain three replicated slices, each with four distinct cell niches. We created “control” by removing two niches from each slice and labeled the original slice as “condition”. This process resulted in three simulated condition pairs (Dataset 1 in Fig. 1b), with the true condition-relevant niches (CRNs) known as ground truth (Fig. 2b).

**Fig. 2.**
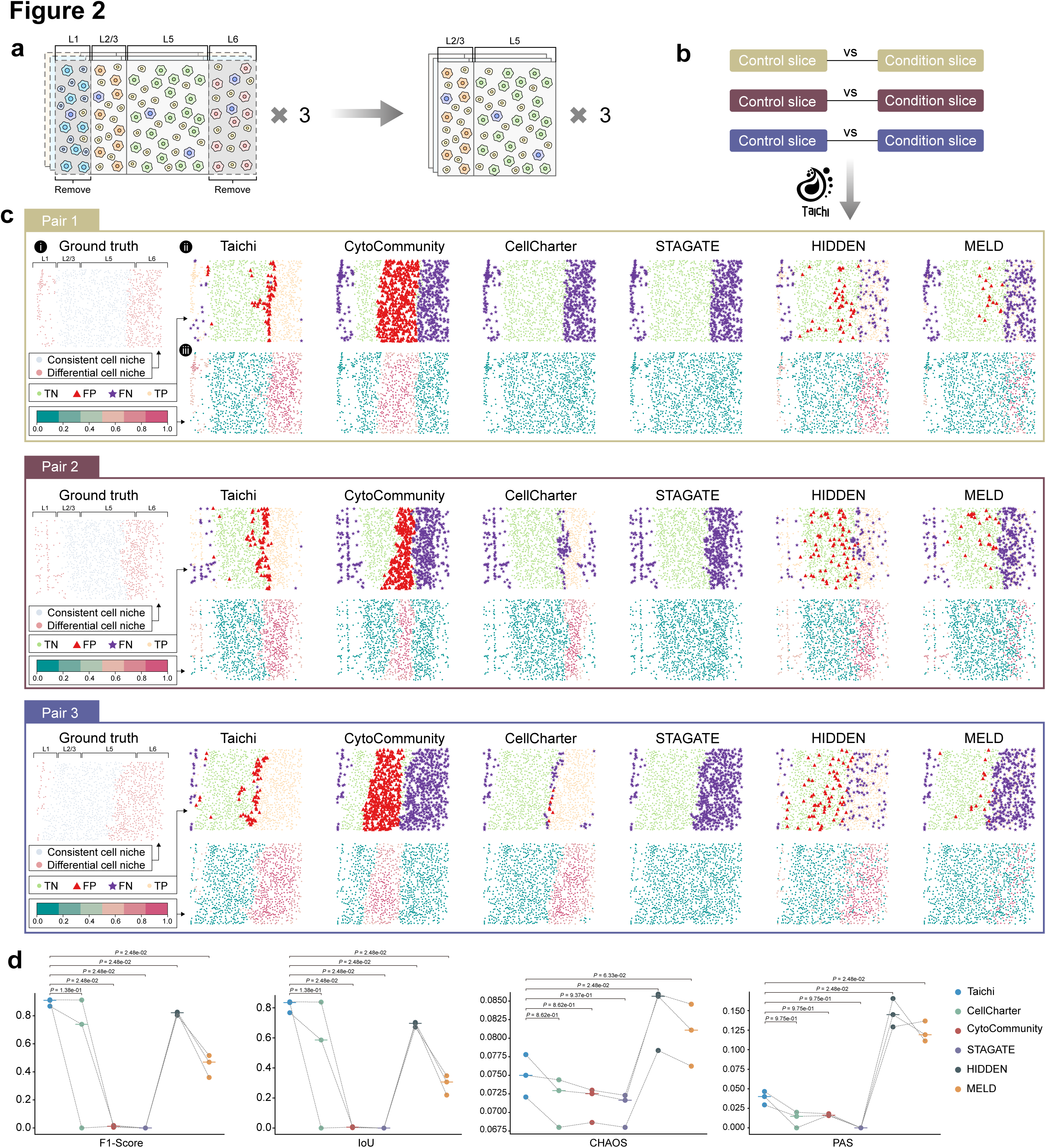
| Simulation pairs obtained by STARmap dataset. **a**, The simulation procedure to obtain the condition and control pairs. Specifically, given that the real datasets contain three replicated slices, each with four layers of expert-annotated cell niches, we generated three condition-control pairs by removing two cell niches, L1 and L6. **b**, In the benchmarking setting for the competing methods, we set each pair as one task and applied each competing method to evaluate method performance. **c**, Benchmarking outcome for each method on three tasks. We take pair 1 as an example, pair 2 and pair 3 are similar: (i) Condition-relevant cell niche (CRN) ground truth. (ii) Results for each method, each cell is labeled as true negative (TN), true positive (TP), false negative (FN), or false positive (FP). (iii) The identified CRN using each method, showing on the spatial map with spatial kernel density. A closer density pattern to the ground truth indicates more similarity to the group truth. **d**, Benchmark results summary for competing methods in three tasks: The metrics represent the identification compared with ground truth: F1-score and IoU, and the metrics indicating the spatial continuity of the result without ground truth: CHAOS, and PAS. P-values are obtained using one-side rank-sum test.

We applied Taichi and other methods to assess their ability to identify these true CRNs (Fig. 2c (i)). We compared the method-identified CRNs (Fig. 2c (iii)) with the true CRNs and visualized the true negative (TN), false positive (FP), false negative (FN), and true positive (TP) in the spatial plot (Fig. 2c (ii)). The fewer red triangles (FP) and purple asterisk markers (FN), the better the method’s performance (Fig. 2c (ii)). Taichi successfully identified most true differential niches in all three condition pairs by pinpointing the missing L1 and L6 niches, which constitute the main difference (Fig. 2c). In contrast, non-spatial methods like MELD and HIDDEN identified cell state-level differences by detecting differential abundance (DA) cells within L1 and L6 regions but missed the spatial information. Spatial clustering-based methods, such as STAGATE, CellCharter, and CytoCommunity, can theoretically consider both cell state and spatial information, but their results were suboptimal. STAGATE failed to identify any true positive (TP) differences in all three condition pairs (Fig. 2c and Supplementary Fig. 2c). CytoCommunity pinpointed incorrect niches compared to the ground truth (Fig. 2c and Supplementary Fig. 2b). CellCharter returned no differences in the first condition pair and missed the niche differences in L1 (Fig. 2c and Supplementary Fig. 2a).

To quantitatively assess the performance of these methods, we used four metrics (see Methods, “Evaluation metrics”): F1-score and IoU for comparing the identified regions with the ground truth, and CHAOS and PAS for evaluating the spatial continuity of the identified cell niches (Fig. 2d). In agreement with the visualization analysis, Taichi outperformed other methods in terms of its ability to identify true differences, as reflected by the highest F1-score and IoU (Fig. 2d). The DA analysis ability of MELD and HIDDEN enabled them to successfully identify individual cells within true differential niches, resulting in moderate F1-score and IoU values, but they lacked spatial continuity (reflected in the highest CHAOS and PAS values, where lower is better) (Fig. 2d). Spatial clustering-based methods obtained good spatial continuity but exhibited diverse performance in terms of F1-score and IoU (Fig. 2d).

### Identifying Different Levels of Perturbations in Complex Tissues

To assess the ability of different methods to identify niches with varying levels of perturbations, we designed a more challenging simulation based on MERFISH data of the mouse hypothalamic preoptic region^41^ (Fig. 3a). The original slice, considered as healthy tissue, contained eight expert-annotated spatial domains: bed nuclei of the stria terminalis (BST), third ventricle (V3), periventricular hypothalamic nucleus (PV), paraventricular nucleus of the thalamus (PVT), medial preoptic nucleus (MPN), columns of the fornix (fx), paraventricular hypothalamic nucleus (PVH), and medial preoptic area (MPA) (Fig. 3a). We simulated eight diseased slices by replacing cells with novel cell types within each domain, creating eight health/disease pairs in each perturbation condition (Fig. 3a, b, see Methods). The complex tissue structure and meticulous simulation design made identifying perturbed niches more challenging compared to the STARmap simulation.

**Fig. 3.**
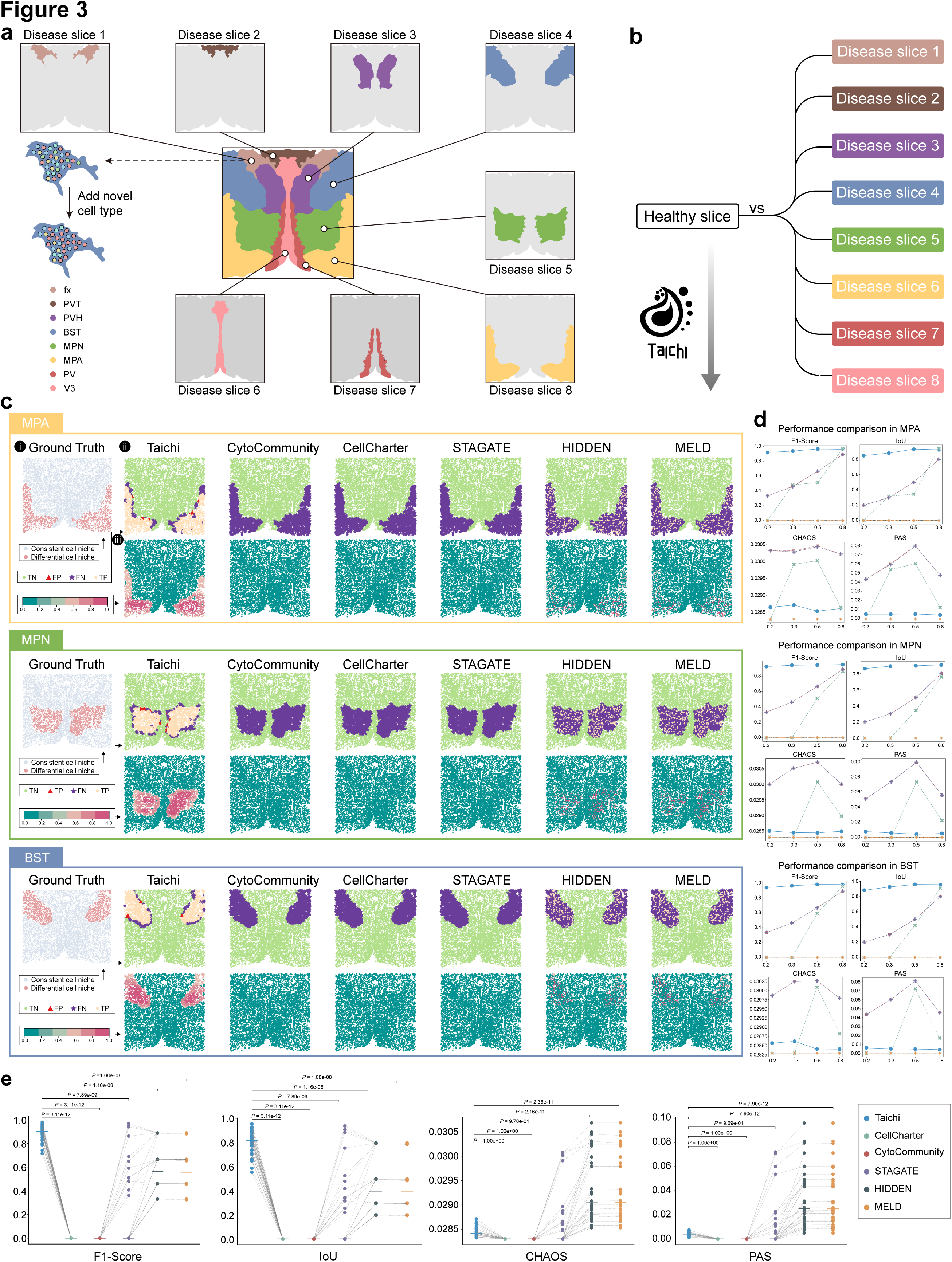
| Simulation pairs obtained by MERFISH dataset. **a,** To benchmark different methods with various levels of perturbations, we designed a simulation based on MERFISH data of the mouse hypothalamic preoptic region. Initially, we considered the original slice as the control slice (healthy slice) and independently perturbed the bed nuclei of the stria terminalis (BST), third ventricle (V3), periventricular hypothalamic nucleus (PV), paraventricular nucleus of the thalamus (PVT), medial preoptic nucleus (MPN), columns of the fornix (fx), paraventricular hypothalamic nucleus (PVH), and medial preoptic area (MPA). This perturbation involved replacing existing cells with our generated “novel cell” at different given fractions, resulting in eight condition slices (diseased slices) at each perturbation fraction (20%, 30%, 50%, 80%) (**b**). **c**, Benchmarking outcome for each method on three tasks. We take the first task (MPA perturbation) as an example, other tasks are similar: (i) Condition-relevant cell niche (CRN). (ii) Results for each method, each cell is labeled as true negative (TN), true positive (TP), false negative (FN), or false positive (FP). (iii) The identified CRN using each method, showing on the spatial map by estimating spatial kernel density. A closer density pattern to the ground truth indicates more similarity to the group truth. **d**, Line plots for the competing method results for different tasks on different perturbation levels in 20%, 30%, 50%, and 80% to show the consistency for the method among different perturbation levels. **e**, Benchmark results summary for competing methods on all 32 tasks. The metrics represent the identification compared with ground truth: F1-score and IoU, and the metrics indicating the spatial continuity of the result without ground truth: CHAOS, and PAS. P-values are obtained using one-side rank-sum test.

We compared the method-identified CRNs with the true CRNs and visualized TN, FP, FN, TP, in the spatial plot (Fig. 3c and Supplementary Fig. 1a-p). A better method performance is indicated by fewer red triangles (FP) and purple asterisk markers (FN). Three representative health/disease pairs are shown as examples (Fig. 3b), with the remaining five pairs demonstrating consistent observations (Supplementary Fig. 1). Single-cell-based methods like MELD and HIDDEN can identify the simulated new individual cells but fail to locate the disease-associated niche area (Fig. 3c and Supplementary Fig. 1a-h), highlighting the importance of integrating spatial information. However, spatial clustering-based methods struggle to locate true CRNs due to their reliance on the identification accuracy of spatial domains (Fig. 3c and Supplementary Fig. 2d-f), a known challenge in complex tissues^45^. Taichi, which does not require pre-defined and discrete spatial clusters, successfully identified true CRNs in all eight datasets (Fig. 3c and Supplementary Fig. 1a-p).

To further evaluate the methods’ performance under different degrees of perturbation, we generated additional diseased tissues by replacing 20%, 30%, 50%, and 80% of cells with new cell types (resulting in 32 disease slices and 1 healthy slice, Dataset 2 in Fig. 1b), with higher new cell type portions leading to easier detection. Taichi consistently achieved high detection power in terms of F1-score and IoU, as well as high spatial continuity measured by CHAOS and PAS, across all perturbation levels (Fig. 3d and Supplementary Fig. 1q). Summarizing the performance of all methods across the eight healthy/diseased pairs, Taichi demonstrated the best performance in terms of F1-score and IoU, along with good spatial continuity, indicating its stable and best performance on complex tissues across different degrees of perturbation (Fig. 3e).

### Cell Type Properties of Disease-relevant Niches

In many diseases, tissue is composed of a mixture of regions with healthy and diseased characteristics. However, current spatial biology studies typically sample slices from diseased tissue and label the whole slice as “disease”, ignoring the existence of healthy-consistent regions within the slice^25–27^. The presence of healthy parts within the tissue can reduce the statistical power or even override disease-relevant insights when performing bioinformatics analysis. Identifying disease-relevant cell niches (which can be considered as purer disease regions) can improve the power of performing in-depth analysis to identify disease processes and therapeutically actionable pathways on those disease-relevant cell niches, rather than on the whole tissue slice.

To demonstrate the applicability of Taichi in such real cases, we collected Slide-seqV2 datasets^46^ of 34 slices of wild-type (WT) mouse kidney and 26 slices of diabetic kidney disease (DKD), containing ∼ 1.5 million cells (beads) in total (Fig. 4a, Dataset 10 in Fig. 1b). The output of Taichi in this dataset is to partition each real DKD slice (expected to be mixture of healthy and disease) into two virtual slices, i.e., slice containing DKD-relevant niche (DRN) and slice containing WT-consistent niche (WCN). The WT-consistent niche is expected to be more similar to WT slices than the DKD-relevant niche. We evaluated such similarity in terms of cell types and cell-cell interactions.

**Fig. 4.**
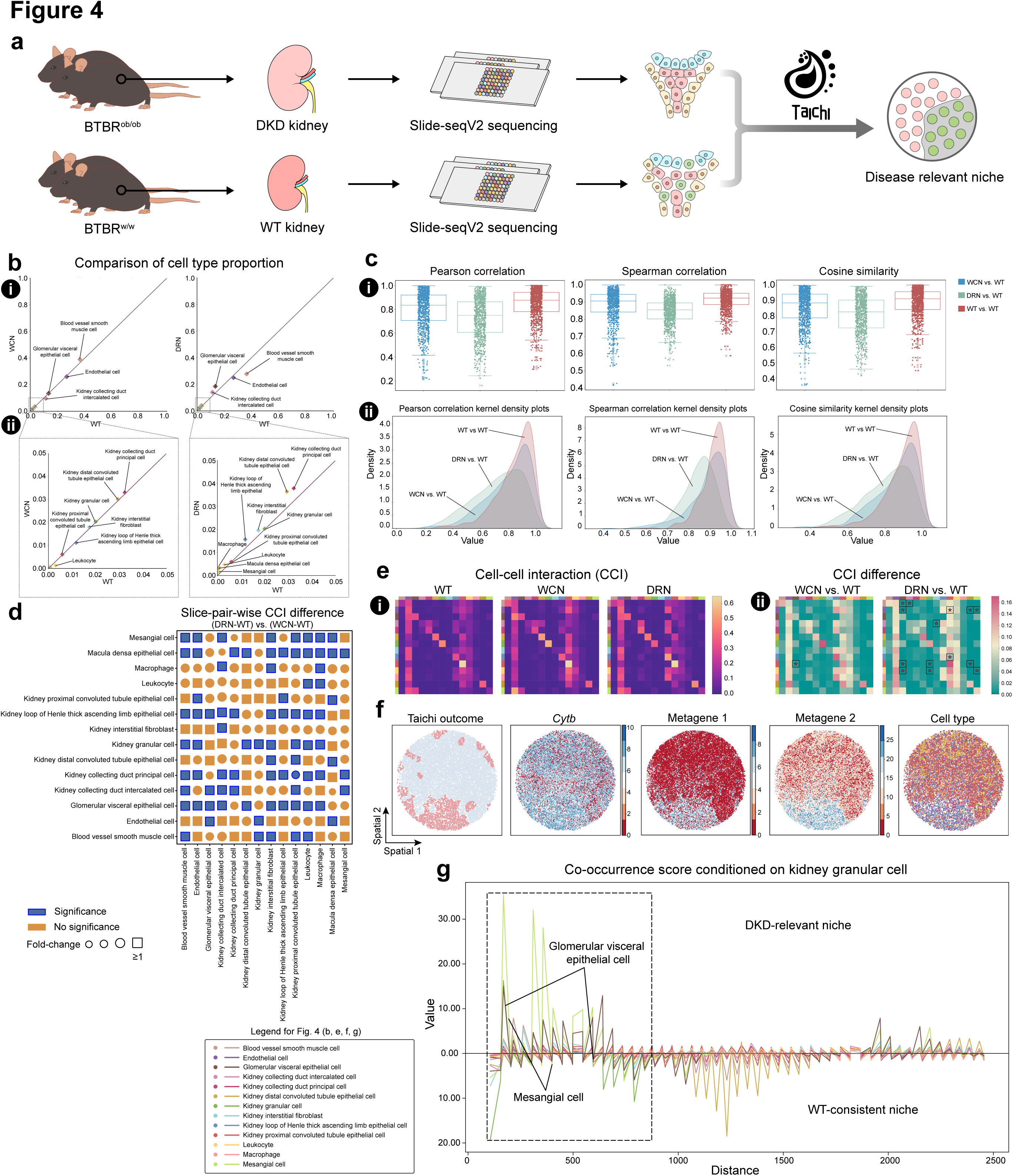
| Cell niche shift in diabetic kidney disease (DKD) compared with wild-type (WT). **a,** Dataset: Slide-seqV2 data from 34 WT (BTBR w/w) and 26 DKD (BTBR ob/ob) mouse kidney slices. Taichi identifies DKD-relevant niches (DRNs) and WT-consistent niches (WCNs). **b**, Cell type proportion comparison between WCN and WT, and DRN and WT. **c**, (i) Slice-pairwise cell type proportion comparisons between WCN vs. WT, DRN vs. WT, and WT vs. WT using Pearson correlation, Spearman correlation, and cosine similarity. (ii) Kernel density comparison for the three metrics, showing point distributions from the boxplots in (i). **d**, Slice-pairwise cell-cell interaction (CCI) differences. Entries indicate interaction differences between cell types. Square entries indicate fold change ratios in DRN to WT/WCN to WT > 1, and blue indicates significant differences (p < 0.05, one-sided rank-sum test). **e**, (i) Median-aggregated CCI matrices for WT, DRN, and WCN slices. (ii) CCI differences for WCN vs. WT and DRN vs. WT. “*” indicates statistical significance (p < 0.05, one-sided rank-sum test). **f**, From left to right: Taichi outcome visualization, DRN-specific gene, DRN-specific metagene 1, DRN-specific metagene 2, and cell type map. **g**, Co-occurrence analysis between kidney granular cells and other cell types in WCN and DRN. Distance indicates the distance to kidney granular cells, and the value represents the co-occurrence proportion of the cell type at this distance.

To analyze the similarity in terms of cell types, we aimed to confirm that the WCN has a more similar cell type proportions to WT compared to the DRN. Using cell type annotations from the original study^46^, we found that the cell type proportions of WCN perfectly correlated with WT for both major (Fig. 4b (i) left) and minor (Fig. 4b (ii) left) cell types. In contrast, the cell type proportions of DRN showed less correlation with WT, as evident from several cell type data points deviating from the y=x function line (Fig. 4b (i, ii) right). To quantify the similarities in cell type proportions among WCN, DRN, and WT more rigorously, we hypothesized that the cell type proportion similarity between a randomly selected slice from WCN and a randomly selected slice from WT should be greater than the proportion similarity between a random slice from DRN and a random slice from WT. We calculated the pairwise proportion similarity between WCN and WT slices (WCN vs. WT) using Pearson correlation coefficient (PCC), resulting in a distribution that closely matched the pairwise slice similarity within the set of WT slices (WT vs. WT) (Fig. 4c (i, ii) left). Both of these distributions were significantly higher than the pairwise slice cell type proportion similarity between DRN and WT slices (DRN vs. WT) (Fig. 4c (i, ii) left). Other metrics, such as Spearman correlation (Fig. 4c (i, ii) middle) and cosine similarity (Fig. 4c (i, ii) right), yielded similar results.

### Spatial Properties of Disease-relevant Niches

For the Slide-seqV2 dataset (Fig. 4a, Dataset 10 in Fig. 1b), we next analyzed the similarity of spatial properties in terms of cell-cell interactions (CCI). The CCI properties in the WT-consistent niche (WCN) within a disease slice are expected to share greater similarity with the WT, whereas the DKD-relevant niche (DRN)’s CCI properties should reflect more disease related biological processes than the disease slice as a whole. We conducted CCI comparison analyses^47^ (see Methods), to demonstrate that the WCN exhibits a CCI pattern more similar to that of WT than does the DRN. We computed the CCI matrix for each slice of WT, DRN, and WCN, assigning a specific CCI matrix to each slice. The WT-CCI matrix displayed a pattern more similar to the WCN-CCI matrix compared to the DRN-CCI matrix (Fig. 4e (i)). Additionally, we quantified the differences in each entry (i.e., the interaction between each pair of cell types) between the WT-CCI matrix and the WCN-CCI matrix (i.e., WCN vs. WT), and between the WT-CCI matrix and the DRN-CCI matrix (i.e., DRN vs. WT) (Fig. 4e (ii)). The analysis revealed that WT and DRN have 14 significantly different CCIs, while WT and WCN exhibit only one significantly different CCI (Fig. 4e (ii)) (see Methods). A notable significant CCI identified by Taichi was between the distal convoluted tubular epithelial cells and the endothelial cells. This interaction is supported by previous studies which have found many common signaling pathways between distal convoluted tubular epithelial cells and endothelial cells, with this crosstalk playing a crucial role in the development and progression of DKD^48^.

To provide a more rigorous and fine-grained analysis, we quantified the interaction differences between DRN and WT, and compared them to the interaction differences between WCN and WT. We computed Cell-Cell Interaction difference matrices (CDMs) by taking the absolute value of the element-wise differences between two CCI matrices, which measure the degree of CCI differences between two conditions. For *n* WCN slices, *m* DRN slices, and *p* WT slices, we obtained *n* × *p* CDMs for WCN vs WT and *m* × *p* CDMs for DRN vs WT, forming two distributions for each CCI. We then performed a one-sided rank-sum test with False Discovery Rate (FDR) control for each CCI to determine which interaction differences in DRN vs WT were significantly larger than those in WCN vs WT. As shown in Fig. 4d, the macula densa (MD) epithelial cells, known indicators of the juxtaglomerular apparatus (JGA)^46^, exhibited the most significant CCI differences (most entries of the row “Macula densa epithelial cell” in Fig. 4d are blue squares). The JGA is known to undergo cellular composition shifts in Diabetic Kidney Disease (DKD)^49,50^.

Since Taichi can partition a single real disease slice (which is a mixture of healthy and disease niches) into two virtual slices—DRN (the true disease slice) and WCN (the healthy slice), to further assess the cell-cell interaction differences between DRN and WCN from a more detailed perspective, we reasoned that additional spatial analysis between the two virtual slices could reveal more meaningful intra-slice differences in cell spatial property. To show the large spatial property difference reflected by co-occurrence patterns between DRN and WCN, we computed co-occurrence probability of the JGA indicator cell type, kidney granular cells, with other cells in the DRN and WCN independently within the same slices (see Methods). The results show that the DRN area exhibits stronger and closer interactions between glomerular-relevant cell types (e.g., mesangial cells, glomerular visceral epithelial cells), indicating the physical proximity of the JGA to the glomerulus and demonstrating an association with the DRN outcomes (Fig. 4g).

### Disease Associated Genes and Metagenes

Gene-expression-centric analysis can provide complementary insights to cell-centric analysis. We utilized the WCN and DRN on the same slice as two distinct spatial domains and identified spatial domain-specific genes and metagenes associated with DKD-relevant niches. Under FDR adjusted p value < 0.05, we identified *Cytb* as a spatial domain-specific gene for the DRN domain, which plays a crucial role in DKD involving reactive oxygen species and exhibits higher expression in diabetic kidneys compared to WT^51^ (Fig. 4f). We also identified metagenes (combinations of multiple individual genes), which are reported to reveal spatial patterns better than any single gene due to the intra-heterogeneity of tissue architecture^29,52^. Using Taichi-identified DRN and WCN (see Methods), we found that these metagenes can more clearly segregate disease-associated regions than using single genes alone (Fig. 4f).

### Identifying Niches Associated with Survival Outcomes in Colorectal Cancer

Taichi can effectively identify key cell niches that define patient groups with different survival outcomes. We demonstrated this using a colorectal cancer (CRC) dataset^53^ obtained from two patient groups: Crohn’s-like reaction (CLR) and diffuse inflammatory infiltration (DII), using CODEX spatial proteomics technology (Fig. 5a, Dataset 11 of Fig. 1b). CLR patients showed significantly longer overall survival compared to DII patients^53^.

**Fig. 5.**
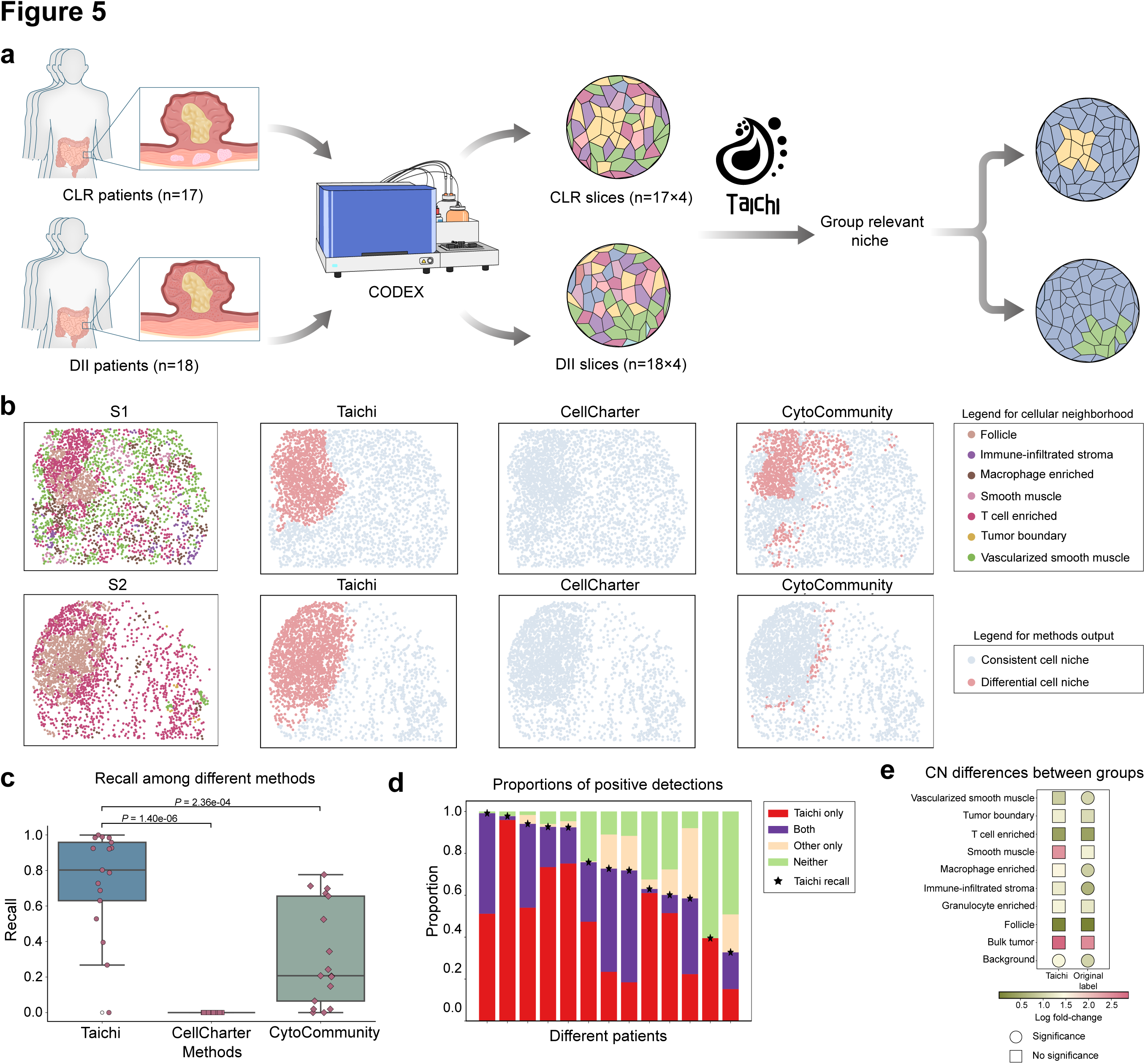
| Survival relevant niches of human Colorectal Cancer (CRC). **a,** Dataset: CODEX data from 17 patients with CLR and 18 with DII. The dataset included 140 tissue regions from 35 patients, with 4 slices per patient, each analyzed using 56 protein markers. Taichi was applied to this multi-slice dataset to identify CLR-relevant niches. **b**, Visualization of CLR-relevant niches identified by Taichi, CytoCommunity, and CellCharter, and corresponding annotations in original study. **c**, Recall comparison among Taichi, CytoCommunity, and CellCharter in detecting tertiary lymphoid structure (TLS) areas in the CLR slice group (TLS represented by the Follicle; hypothesis test conducted by one-sided rank-sum test). Taichi’s recall is significantly higher than CytoCommunity and CellCharter (p < 0.05). **d**, TLS identification among patients, categorized by: identified only by Taichi, identified only by other two methods (CellCharter, CytoCommunity), identified by all three methods, and not identified by any method. **e,** Differential abundance of CNs between CLR and DII groups under two grouping strategies (original label indicates original slice-level annotated group, and Taichi indicates grouping by Taichi-identified CLR-relevant and DII-relevant slices). Squares indicate significant differences (p < 0.05 by rank-sum test), with magnitude representing the log fold change.

Tertiary lymphoid structures (TLS) have been shown to have high biological and clinical relevance in CRC and serve as key differences between CLR and DII groups^54,55^. Quantifying whether the method-identified differential niches can cover TLS regions (i.e., known differential niches) can evaluate the performance of different methods. The originally annotated TLS structure (labeled as “Follicle”) overlapped more with Taichi than with two recent spatial methods, CellCharter and CytoCommunity (Fig. 5b). Quantitative analysis across all patient slices showed that Taichi had the best detection of TLS structures compared to other methods (Fig. 5c, d).

In the original study, each slice was labeled as a whole according to its patient group, which is noisy. Taichi partitioned each slice into two virtual slices with more fine-grained and accurate labels than originally provided. We assume that the refined labels obtained by Taichi can identify more meaningful differences than the original noisy labels. Comparing the abundance of cellular neighborhoods (CN, annotated by the original paper) between originally labeled CLR/DII slice groups revealed that 6 out of 10 CNs were significantly different in abundance (p-value < 0.05). Using Taichi-refined labels, we found additional significantly different CNs, such as “Macrophage enriched” and “Immune-infiltrated stroma” (Fig. 5e). The Macrophage-enriched CN, characterized by an increased ratio of CD163+/CD68+ macrophages and a lack of single CD68+ macrophages, corresponds to previous studies showing that an increased ratio of CD163+/CD68+ in the tumor invasive front was positively correlated with shorter CRC relapse-free survival and overall survival time^56^, echoing the shorter survival property of the DII group.

### Improving Clinical Phenotyping on Head and Neck Cancer and Colorectal Cancer

In disease research with complex experimental designs, one sample slice may be annotated with multiple kinds of labels to reflect different aspects of the patient’s situation or treatment outcome. In previous sections, we demonstrated Taichi’s capability to capture and identify condition-relevant niches from the originally annotated coarse-grained slice-level labels derived from patient phenotypes. Here, we assumed that the distribution of condition-relevant niches themselves can also act as indicators of different aspects of the patient’s situation or treatment beyond the original label in the corresponding slice. This hypothesis is based on the observation that the pre-defined slice-level phenotype labels are coarse-grained and do not account for intra-slice heterogeneity. Meanwhile, variations in the distribution of condition-relevant niches may reveal such heterogeneity, as they filter out niches irrelevant to the phenotype.

To show the potential of Taichi for clinical applications with the pre-defined coarse-grained labels, we applied Taichi to 3 large-scale and multi-slice spatial datasets containing 2,934,175 cells from 103 slices in total, with slice-level phenotypic annotations^57^ (Dataset 12-15 in Fig. 1b). We expected to see that Taichi could utilize the distribution of condition-relevant niches to determine more fine-grained clinical labels (Fig. 6a). We applied Taichi to a multi-slice dataset and utilized the area of the condition-relevant niches as a discriminative property to reflect intra-group heterogeneity within the corresponding slices. Specifically, we assigned a score to each slice within the original condition slice groups based on the area of the condition-relevant niches (see Methods) and used this score for further analysis.

**Fig. 6.**
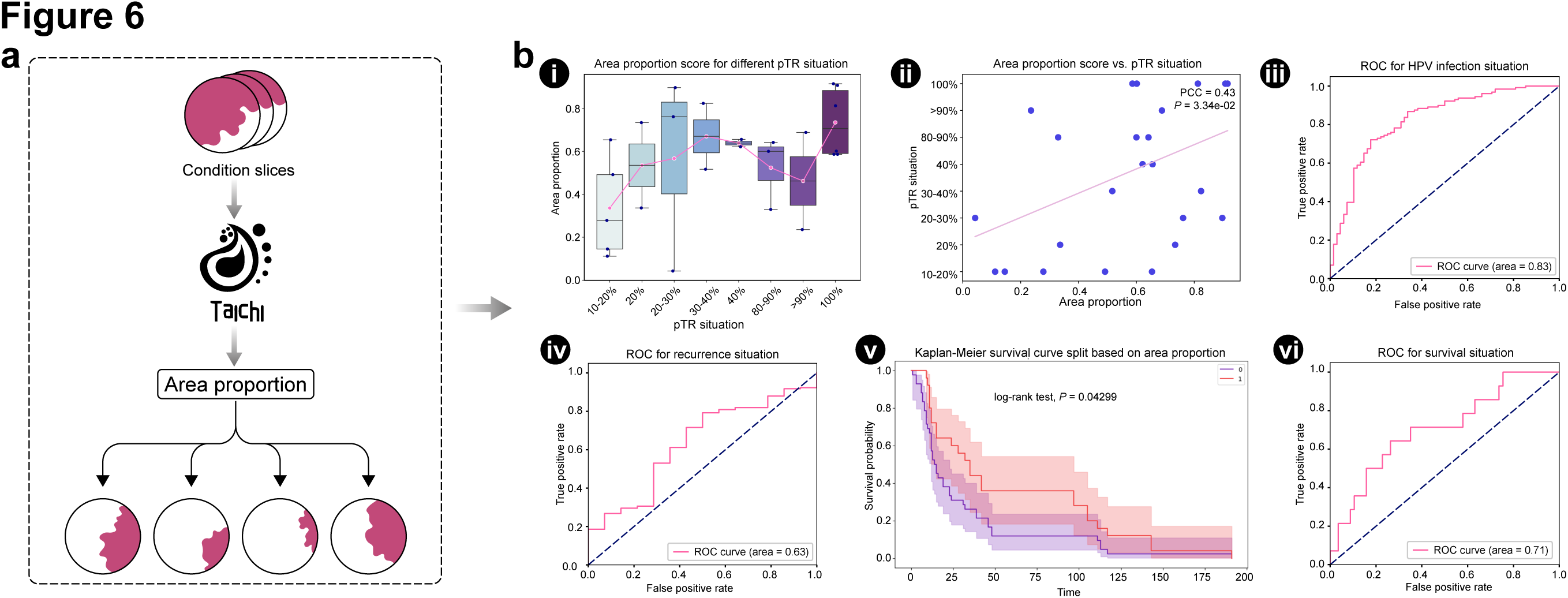
| Fine-grained patient stratification using Taichi-identified niches in various human cancers. **a**, Workflow illustrating the use of Taichi-identified niches to determine more fine-grained patient properties. **b**, Fine-grained partition results on different tumor CODEX datasets: (i) In the DFCI-HNC dataset, Taichi higher-pCR-relevant niches area fraction box plot shows the association of the area fraction score with continuous pCR scores across several thresholds (10-20, 20-30, 30-40, 40, 80-90, >90, 100). (ii) In the DFCI-HNC dataset, the correlation between condition-relevant niches area fraction and continuous pCR scores across several thresholds (10-20, 20-30, 30-40, 40, 80-90, >90, 100) is analyzed using Pearson correlation. (iii) The ROC-AUC of HPV infection status is determined by Taichi condition-relevant niches area fraction of the evidence-of-disease condition as a score in the UPMC-HNC dataset. (iv) The ROC-AUC of recurrence status is determined by Taichi condition-relevant niches area fraction of the evidence-of-disease condition as a score in the UPMC-HNC dataset. (v) The Kaplan-Meier survival curve of two groups (1 indicates high risk and 0 indicates low risk), partitioned by the mean of deceased-relevant niches area, shows statistical significance (log-rank test, p-value = 0.04299) in the Stanford-CRC dataset. (vi) The ROC-AUC of deceased/recurred status within 60 months is determined by Taichi condition-relevant niches area fraction of the deceased condition as a score in the Stanford-CRC dataset.

The first analysis utilized the condition-relevant niche area fraction derived from Taichi with the original Disease-Evidence (DE) condition label of UPMC-HNC (Head and Neck Cancer) slices. Specifically, we applied Taichi to the UPMC-HNC dataset, which consists of 111 DE slices and 197 non-DE slices, with DE as the condition label. Since the Taichi outcome’s condition-relevant niche is associated with the DE property of HNC, we assume that the niche area fraction on the slice may indicate the subtype and situation of the HNC sample. Thus, we utilize the area fraction score as an indicator of both HPV infection and non-recurrence in the DE groups. As shown in Fig. 6b, the niche fraction demonstrates the capability to discriminate between HPV-infected and non-recurrent samples within the original DE groups, distinctions that were not possible with the previously annotated single label (Fig. 6b (iii, iv)) (see Methods).

Furthermore, following the same procedure used on the UPMC-HNC dataset, we applied Taichi and obtained the condition-relevant niche area fraction score from another dataset—Stanford-CRC slices labeled with the deceased condition (71 slices) and control survival (292 slices). The deceased-relevant niches contain the cell spatial properties determining the disease risk of each sample, and thus this score can be regarded as an indicator of recurrence or death within 60 months for each sample, as depicted in (Fig. 6b (vi)). Since a larger deceased-related niche reflects a higher risk of recurrence or death within 60 months, we utilized the mean niche area value to partition patients into higher-risk (larger area fraction score) and lower-risk (smaller area fraction score) groups. The motivation here is that a larger deceased-related niche corresponds to a higher risk of death and thus indicates shorter survival. This partition is evaluated based on survival length (see Methods), as shown in (Fig. 6b (v)).

In the third dataset, DFCI-HNC, which focuses on high primary tumor response to treatment (pTR > 10%) (25 slices) as the condition and low pTR < 10% (33 slices) as control, the niche area fraction is calculated under the supervision of this pre-defined coarse-grained label in the same manner. Different from the previous two datasets, the additional property of the given label belongs to the same category but is more fine-grained. Since the high pTR-relevant niches are associated with the pTR condition on the slice, the abundance of these niches can represent the specific degree of the pTR. Thus, we subsequently utilized the resulting niche area fraction as a score to assess more fine-grained pTR situations. The results show that this approach effectively captures the continuous variation in pTR without the need for additional information, as illustrated in (Fig. 6b (i, ii)). These findings demonstrate that the Taichi-captured condition-relevant area contains more fine-grained and diverse information beyond the pre-defined labels, supporting the potential clinical application of Taichi.

### Revealing Complex interaction related to Triple Negative Breast Cancer Immunotherapy Response

In the previous section, we explored the potential of Taichi for disease diagnosis through the analysis of condition-relevant niche area fractions. This application highlights Taichi’s capacity for fine-grained clinical classification. In this section, we aim to demonstrate how Taichi can also assist in identifying key biomarkers for the immunotherapy response on Triple Negative Breast Cancer (TNBC) of pre-treatment sample incorporated with Taichi and tensor decomposition (Fig. 7a).

**Fig. 7.**
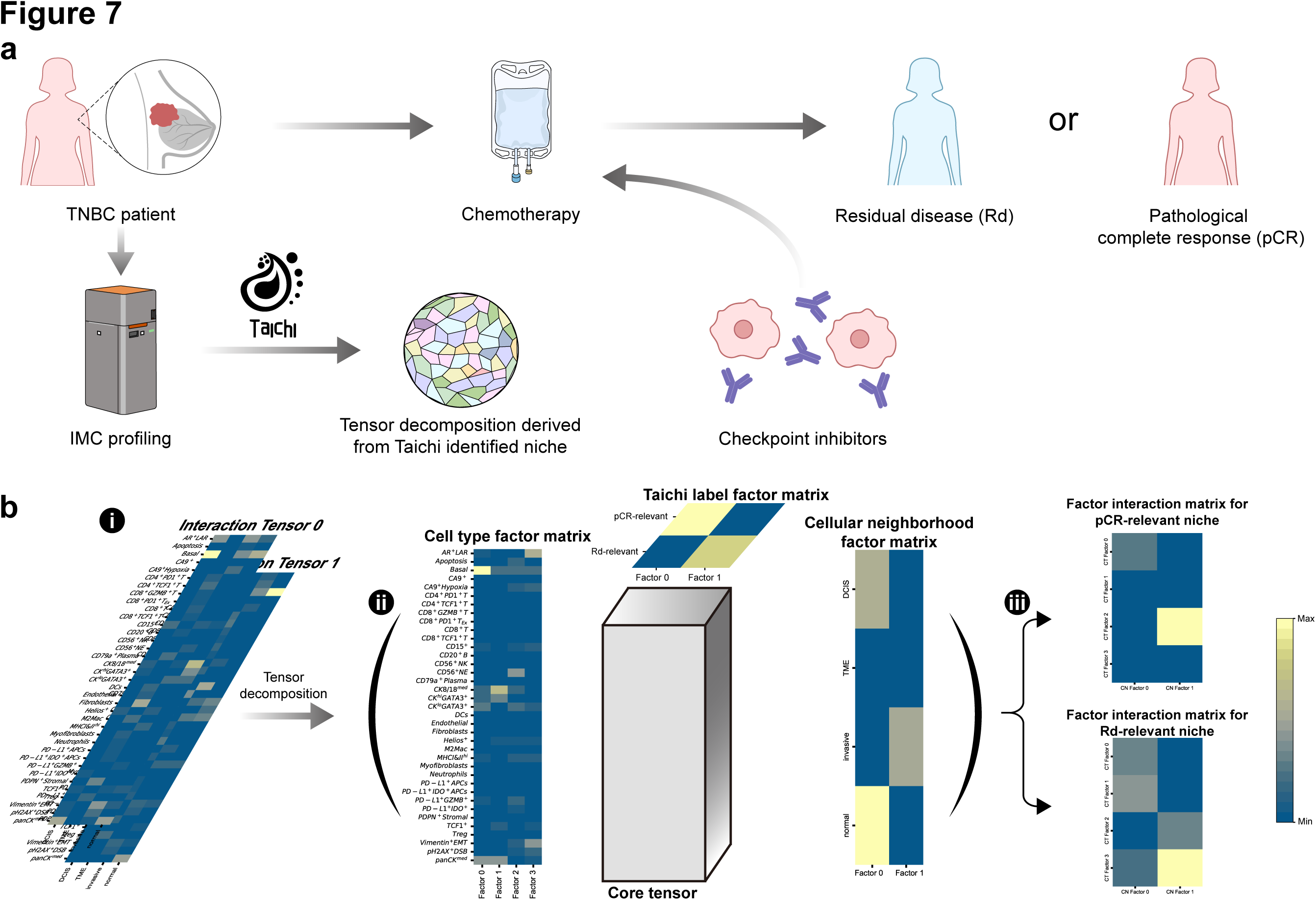
| Immunotherapy response biomarker identification in human Triple-Negative Breast Cancer (TNBC) using Taichi outcome niches. **a**, Workflow of the TNBC IMC data source: The multi-slice TNBC spatial proteomics dataset is obtained from patients before immunotherapy and chemotherapy and is labeled according to the post-treatment response—either pCR (Pathological Complete Response) or Rd (Residual Disease). **b**, (i) Input for the Tucker decomposition includes a stack of two cell type proportion matrices on different cell neighborhoods, one for cells belonging to Taichi-identified pCR-relevant niches and the other for Rd-relevant niches. (ii) The decomposition result includes the cell type factor matrix, cell neighborhood factor matrix, and Taichi niches label factor matrix with the core tensor. (iii) The factor interaction matrix corresponds to the two Taichi niche labels (pCR-relevant and Rd-relevant) for the cell type and cell neighborhood factors. The interaction matrices are obtained by the multiplication of the core tensor with the cell type and cell neighborhood factor matrix tensor, and each tensor layer (matrix) is classified as pCR/Rd-relevant depending on the factor value in the Taichi niche label factor matrix.

We applied Taichi to a large-scale TNBC Imaging Mass Cytometry (IMC) dataset^27^ (Dataset 15 in Fig. 1b) annotated with immune therapy responses (pathological complete response (pCR) or not), focusing on the baseline sample group (before treatment). This approach is particularly valuable for identifying biomarkers related to treatment response before the commencement of therapy, as opposed to during treatment. Utilizing the Taichi-defined condition-relevant and control-relevant niches under the pCR condition setting, we constructed a tensor representing the fraction of each cell type (CT) within the originally annotated cell neighborhood (CN) for each Taichi niche, as depicted in (Fig. 7b (i)). This tensor was decomposed into two distinct factor interaction compartments for the two Taichi niches (Fig. 7b (iii)) (see Methods). Each factor represents a combination of different cell types and their microenvironment (Fig. 7b (ii)). This compartmentalization reveals the interplay between latent factors of cell types and cell neighborhood, a nuanced analysis made possible only with Taichi’s partitioning of the slice. For instance, the interaction module in the pCR-relevant niche shows significant activation between CN factor 2 and CT factor 3 (Fig. 7b (iii)), which includes CD56+ Neuroendocrine (NE) cells within an invasive cell neighborhood. This was confirmed by previous studies that indicated higher expression of CD56+ NE cells correlates with a higher likelihood of pCR in immunotherapy^58^.

### Scalability and Speed

Scalability and Speed We evaluated Taichi’s scalability and speed in two common spatial omics research scenarios: S1, involving a large number of slices with a moderate cell count per slice, and S2, featuring a large number of cells per slice with a moderate number of slices. S1 is prevalent in disease research across different patients, while S2 is typical in recent high-throughput, single-cell resolved imaging-based studies. S2 challenges spatial clustering-based methods due to the high computational cost of maintaining large graphs^45^.

To compare Taichi with other spatial methods (STAGATE, CellCharter, and CytoCommunity), we simulated large datasets mimicking S1 and S2 (see Methods). Fig. 8a reports the results, considering running times exceeding 8 hours as unavailable. Taichi demonstrates the most efficient and scalable performance in both scenarios. In S1, most methods perform adequately, except for the GNN-based STAGATE, which is limited by GPU memory. Taichi exhibits a nearly linear increase in running time when scaling to more slices, requiring only 1 hour for 5 million cells across 1000 slices. In S2, only CellCharter and Taichi can handle the large-scale single-slice condition, highlighting the advantage of methods not heavily reliant on spatial graph modeling. Taichi outperforms CellCharter with an 18-fold improvement in efficiency, handling an extremely large dataset of 16 million cell in 750s.

**Fig. 8.**
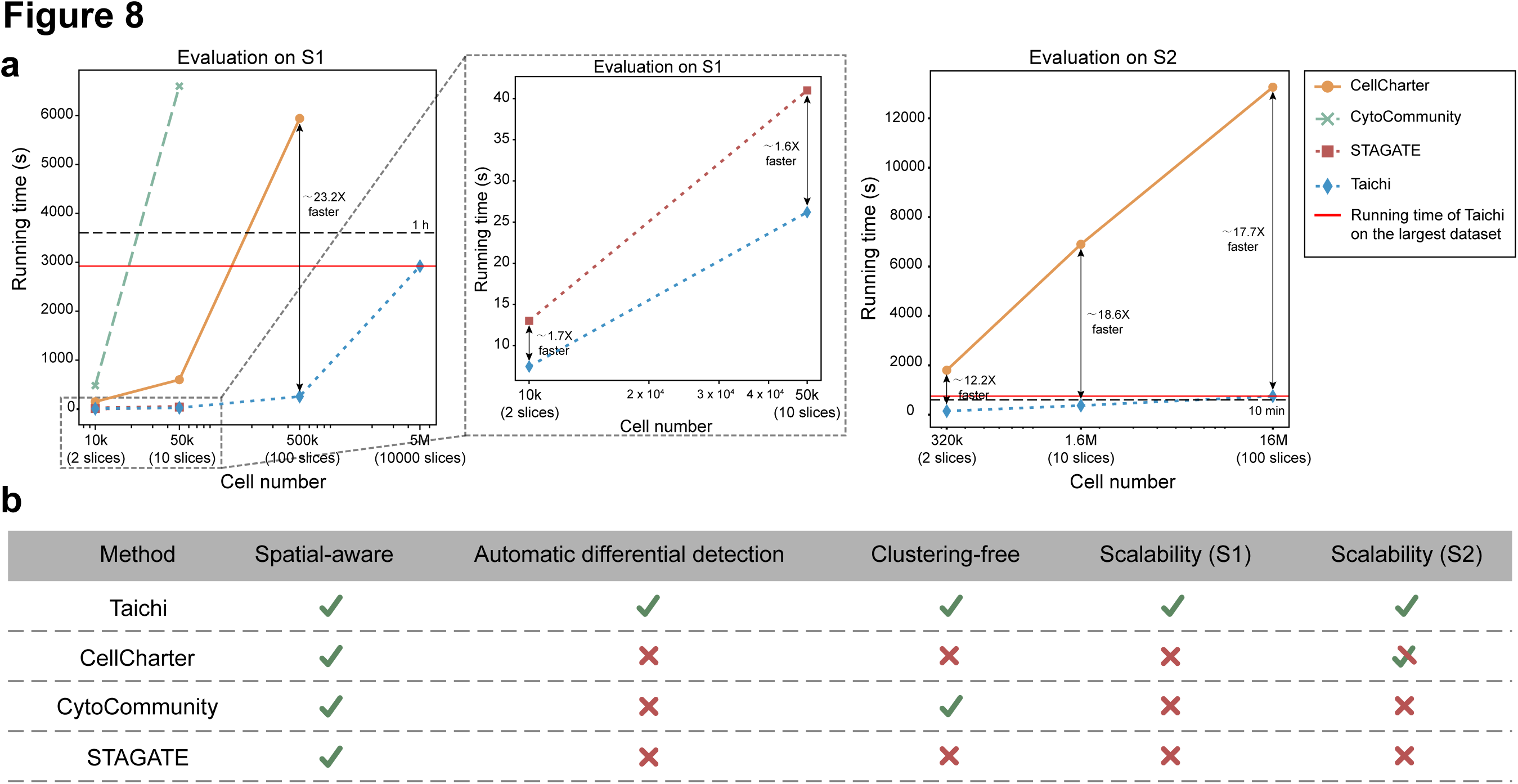
| Scalability benchmarking and functional comparison of methods. **a**, Running time comparison between different methods in two scenarios: S1 (large number of slices with a small number of cells per slice) and S2 (small number of slices with a large number of cells per slice). Only the methods that completed the analysis without running out of memory (OOM) and within 8 hours are shown in the figure. **b**, Functional comparison between competing methods. “Spatial-aware” indicates that the model utilizes spatial information from the input slices. “Automatic differential detection” indicates that the method can directly output the target niches without manual intervention. “Clustering-free” indicates that the method does not depend on spatial clustering or predefined spatial domains. “Scalability” indicates that the method can scale well in S1 and S2 scenarios.

## Discussion

In this study, we introduced Taichi, a highly efficient and scalable computational method for condition-relevant cell niche (CRN) analysis in large-scale spatial omics datasets. Taichi addresses the limitations of current disease studies that often overlook intra-slice niche heterogeneity and fail to identify condition-relevant cell niches. Taichi also eliminates the need for tissue prior knowledge and pre-defined discrete spatial clustering. The name “Taichi” draws inspiration from the philosophical concept of “Taiji” (Yin and Yang), which emphasizes the coexistence and interplay of opposing forces or elements. Just as Taiji represents the balance and harmony between Yin and Yang, Taichi recognizes the presence of both normal and disease-associated cell niches within diseased tissues. By applying this principle to the analysis of spatial omics data, Taichi enables the identification and separation of distinct niche types, providing valuable insights into the underlying mechanisms of disease.

One of the key advantages of Taichi is its ability to delineate major shifts in cell niches between healthy and disease conditions. In our analysis of a mouse model of diabetic kidney disease (DKD), Taichi accurately identified the key differential niche-associated metagenes, providing valuable insights into the molecular mechanisms underlying the disease. This demonstrates the potential of Taichi to uncover spatially resolved insights into disease-relevant niches and advance our understanding of complex biological processes. Furthermore, Taichi’s application to colorectal cancer (CRC) datasets revealed its ability to identify key differential niches between patient groups, resulting in significantly different survival outcomes. This finding highlights the clinical relevance of Taichi, as the identified differential niches could serve as potential biomarkers or therapeutic targets. Moreover, the differential cellular neighborhoods identified by Taichi were more meaningful than the original labels, suggesting that Taichi can uncover novel insights that may be overlooked by traditional approaches.

Taichi demonstrates significant potential in improving clinical phenotyping for head and neck cancer (HNC) and colorectal cancer by leveraging the distribution of condition-relevant niches to uncover intra-slice heterogeneity and provide fine-grained insights beyond original coarse-grained labels. In various datasets, Taichi’s condition-relevant niche area fractions effectively discriminated between clinically relevant subgroups, such as HPV-infected and non-recurrent samples, higher-risk and lower-risk patients, and continuous variations in primary tumor response. These findings highlight Taichi’s ability to uncover valuable information beyond pre-defined labels.

Furthermore, Taichi’s capabilities extend to identifying key biomarkers for immunotherapy response in Triple Negative Breast Cancer (TNBC). By applying Taichi to a large-scale TNBC Imaging Mass Cytometry dataset and performing tensor decomposition, distinct factor interaction compartments were revealed for condition-relevant and control-relevant niches. This compartmentalization allowed for the analysis of the interplay between latent factors of cell types and cell neighborhoods, leading to the identification of significant activation between specific factors, such as CD56+ Neuroendocrine cells within an invasive cellular neighborhood, which aligns with previous studies. These findings demonstrate Taichi’s potential in identifying key biomarkers for treatment response.

The speed and scalability of Taichi are another significant advantage, as demonstrated by our benchmarking results. As spatial omics data become increasingly available through various consortia efforts, the ability to efficiently analyze large-scale datasets is crucial. Taichi’s scalability makes it well-suited for such analyses, enabling researchers to explore complex biological questions across extensive multi-slice and multi-condition datasets. We have compared Taichi and other spatial methods in Fig. 8b.

There are several points that can be improved in future studies. Firstly, Taichi can be enhanced by integration with a large transcriptomics language model pre-trained on extensive corpora beyond the input data. This adaptation facilitates a deeper understanding of the niche, thereby improving Taichi’s performance. Additionally, adapting Taichi to analyze continuous conditions, such as developmental stages, enables the discovery of condition-relevant niches that vary over time. This extension broadens Taichi’s applicability, allowing it to address a more diverse range of applications in future spatial omics research.

## Methods

### Notation

Suppose the input data contains of *K* spatial omics slices, including both condition and control slices. Each slice’s (e.g., *S_k_*) has a binary label **y*_sk_*, where 1 indicates a condition slice and 0 indicates a control slice. Within each slice (e.g., *S_k_*), there are *n_k_* cells (spots or beads for certain spatial technologies, we used “cell” for convenience), and each cell *i* is annotated with a cell type *c_i_* in {*C*_1_, …, *C_M_*}, derived from either expert annotations or post-hoc methods, such as single cell clustering^59^. Additionally, each cell is assigned a spatial coordinate noted as *x_i_*. The pre-defined input label *y_i_* for each cell in the *slice_k_* is equal to **y*_sk_*.

### Spatial Co-embedding

Multi-range cEll coNtext DEciphereR (MENDER) ^39^ is a non-parametric approach designed to extract features from spatial omics data niche, emphasizing the spatial distribution of cell types/states. It leverages multi-resolution cell neighborhood statistics, to capture the local cellular neighborhood information across multiple scales.

For each slice *slice_k_*, consider the spatial location matrix *X*, for each cell has coordinates denoted by *x_i_*, we define the distance between any two cells *x_i_* and *x_j_* by the Euclidean distance:

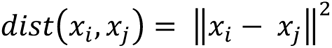

After the distance measurement definition, two hyperparameters *S* and *R*, where *S* controls the number of scales to capture the cellular neighborhood, *R* defines the radius for each scale range, are defined. Then, for each scale *s* in {1, …, *S*} and each cell type *C_m_*, we calculate the score for cell *i* on the *slice_k_* as:

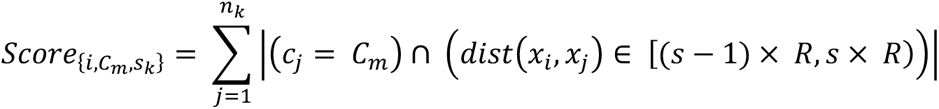

After obtaining the score for each cell type in all scales for cell/spot *i*, we can construct the embedding vector for each cell type *C_m_* for cell *i*, *M*_{*i,cm*}_, by concatenating the score for all cell types in all scales:

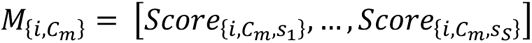

Consequently, the embedding for each cell, denoted by *i*, is derived by aggregating the feature vectors *M*_{*i,cm*}_ for all cell types *C_m_*,

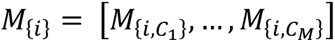

This procedure is applied to all input slices, maintaining consistency in cell annotations across different slices to mitigate batch effects and facilitate an efficient and biologically-consistent co-embedding for multiple slices (“Batch effects”). This embedding can be effectively utilized as an input feature in subsequent analyses.

### Batch effects

Taichi effectively handles batch effects through its use of the MENDER representation, which requires aligned cell type annotations across slices as input. If the cell types across slices are expert-annotated by the original publication and considered reliable, Taichi can directly utilize these annotations, leveraging expert knowledge and ensuring consistency in cell type definitions. However, when expert-annotated cell types are unavailable or unreliable, Taichi employs a two-step process. First, existing batch effect removal tools, such as Harmony^60^, are applied to minimize technical variations between slices and align the data from different batches. Then, single-cell clustering^61^ is performed on the aligned data to identify cell types or cell states across slices. The resulting aligned cell clustering labels are used as input for the MENDER representation, ensuring consistent cell type definitions across slices and mitigating the impact of batch effects. This flexible approach allows Taichi to leverage expert knowledge when available and reliable, while also providing a robust alternative that utilizes existing bioinformatics tools for batch effect removal and cell type identification when needed.

### Label refinement

After extracting the MENDER embedding, we aim to use the control and condition slice labels to identify cell niches that are specifically abundant under conditions. For this purpose and inspired by the noisy label problem in previous studies^42^, we performed label refinement that offers the scalability required for large-scale multi-slice experiments. With the aforementioned MENDER embedding *M_i_* for each cell niche, we proceed with the following procedure:

Firstly, we train the logistic regression model given the MENDER embedding for each cell niche as input feature and the original label for the corresponding slice-level annotation *y_i_* as the label. After determining the model coefficients β_0_, β_1_, …, β*_n_*. that best fit the data, these coefficients are utilized to compute the estimated probabilities. These probabilities serve as scores representing each cell.

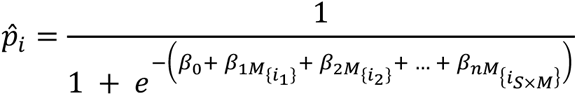

where 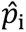 is the estimated probability that *M_i_* belongs to the condition class, and we will utilize this score in the following procedure.

Then, we apply K-means clustering on the score obtained in last procedure to select the higher condition relevant cell niches. The rationale is that not every niche within a condition-labeled slice is intrinsically associated with the specific condition; for example, not all niches in tumor tissue are relevant to tumor. We implement K-means clustering on the logistic regression-derived scores to bifurcate them into two distinct clusters, based on the hypothesis that true positive instances will manifest higher probability scores than false positive instances. Accordingly, we perform K-means clustering with *k* = 2 on to categorize the cells into two groups. The cluster with the higher mean value of 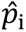 is annotated as the refined positive (condition) label. The remaining instances are annotated as the refined negative (control) label in original condition sample group. We denote the refined label as 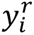, where 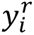 indicates the label assigned to each instance after the refinement process.

### Graph diffusion

After getting the refined labels 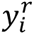, we employ graph heat diffusion to spatially propagate the refined label signals. This process aims to transform the isolated labels, derived from the embedding space, into continuous labels within the original tissue space. The objective is to achieve a more spatially continuous labeling, ensuring that the labels accurately reflect the underlying spatial relationships and patterns. We independently execute the same procedure for each slice within the dataset. Given the featured graph *G_i_* constructed from each slice’s spatial location matrix with nearest neighbors using *squidpy.gr.spatial_neighbors*, we have vertices *V* and edges *E* and node features represented by each cell/spot’s corresponding refined label 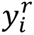. We then conduct heat diffusion for the refined labels among the graph. This process reduces isolated areas and smoothens the transitions between connected communities of the same label. Connected positive areas are maintained by different heat sources, while isolated positive (condition) areas are influenced by the neighboring negative (control) areas.

Let *Y^r^* ∈ *R*^n.^ be the node feature matrix. The procedure for applying heat diffusion to these feature vectors is as follows: First, we construct the Laplacian matrix *L* where *L* = *D* − *A*. Here, *A* is the adjacency matrix of the graph, and *D* is the diagonal degree matrix of *A*. With the Laplacian matrix, we can then construct the heat diffusion process over time *t* for the feature vectors governed by the differential equation:

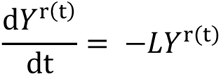

where *Y*^r^ represents the state of the feature vectors at time *t*.

The solution to this differential equation can be expressed using the matrix exponential:

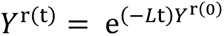

where *Y*^r(0)^ the initial state of the feature vectors, and *Y*^r^ ^(t)^ denotes the features diffused across the graph after time *t*.

We utilize the results at the 25th time point of the diffusion process. Following the diffusion procedure, each cell niche is assigned a final score 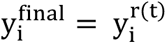, which indicates its relevance to the condition. This score is then used for the final tissue partitioning, where each cell niche is categorized as either condition-relevant or control-relevant on the original slice. We apply this procedure to every pre-defined condition slice in the dataset to determine the final detection outcomes of Taichi.

## Simulation data for benchmarking

### Condition pairs simulated from STARmap datasets

To create a controlled environment for evaluating the performance of different methods, we simulated condition pairs using the STARmap dataset of the mouse brain cortex. We established three condition pairs by removing the most lateral sections (left and right) from each slice. The original slice was labeled as the “condition”, while the modified slice without the lateral sections was labeled as the “control”. The primary difference between the control and condition groups is the absence of the two lateral brain layers. The performance of each method was assessed based on its ability to identify these condition-specific domains within the slices.

### Condition pairs simulated from MERFISH datasets

To further evaluate the robustness of the models under more complex and variable conditions, we utilized a MERFISH mouse brain dataset with more complex tissue structures. We introduced variability by randomly converting different proportions (20%, 30%, 50%, and 80%) of cell types within eight pre-defined spatial domains into a novel cell type. To ensure comparability with other gene expression modification techniques, we also adjusted the gene expression levels of the selected cells. Specifically, we increased the expression of 30 genes by two-fold, another 30 genes by 1.5-fold, and decreased the remaining 95 genes by 0.5-fold. This procedure was implemented across the eight annotated spatial domains at the four perturbation levels, generating a total of 32 distinct tasks. This simulation allows for a comprehensive evaluation of each model’s capability to discern perturbed spatial domains under diverse conditions, assessing their stability and robustness against various degrees of perturbations.

### Evaluation metrics

To assess the methods performance in condition relevant niche detection, several metrics are employed, including Precision, Recall, F1-Score, and Intersection over Union (IoU) score. Formally, the F1-Score is denoted by:

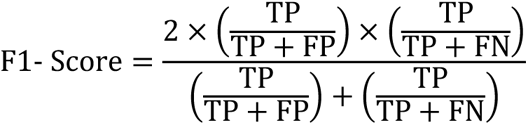

Where the TP (True Positive), FP (False Positive), TN (True Negative) and FN (False Negative) indicates four model prediction situations.

The IoU score, commonly used in object detection scenarios, quantifies the overlap between the predicted and actual bounding boxes. It is calculated as the ratio of the area of overlap to the area of union between the predicted bounding box and the ground truth. This metric ranges from 0 to 1, with 1 indicating a perfect match, and serves as a robust indicator of detection accuracy.

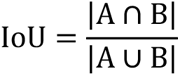

Where the A indicates the area our model predication and B indicates the ground truth area.

Additionally, we incorporate the PAS and CHAOS scores, previously utilized in spatial domain analysis^62^, to evaluate the spatial continuity of the result regions. Lower values indicating higher spatial continuity of predictions.

The PAS score is a measure used to assess the spatial homogeneity of domain identification methods in spatial transcriptomics. A decreased PAS score indicates a higher level of continuity within the identified spatial domains, suggesting greater spatial homogeneity. The score is calculated by determining the proportion of cells whose spatial domain assignment is inconsistent with at least six of their ten adjacent cells.

The CHAOS score is a performance metric that evaluates the spatial continuity of mass spectrometry imaging and spatial transcriptomics. A reduced CHAOS score implies enhanced continuity in the detection of spatial domains. To apply the CHAOS score for assessing the performance of each niche identification task, we begin by constructing a one-hop nearest neighbor (NN) graph for each slice. In this graph, each cell is connected to its nearest neighboring cell based on the minimum Euclidean distance within the slice’s physical space. Given a niche *k* and the Euclidean distance *d* between *cell_i_* and *cell_j_* in the physical space, we calculate *w_kij_* as *d_ij_* if *cell_i_* is connected to *cell_j_*, and set it to 0 otherwise. Finally, we obtain the CHAOS score for the slice by aggregating the scores across all niches and cells.

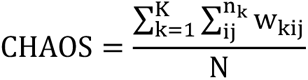

### Competing methods

In this study, we compare three main categories of methods for identifying condition-relevant differences in spatial transcriptomics data.

The first category contains spatial co-clustering methods, including STAGATE^34^ and CellCharter^22^. They are directly applicable to spatial transcriptomics data. These methods extract embeddings for each cell niche and perform clustering to identify discrete niche clusters (also known as spatial domains). The differences in the niche labels between conditions serve as indicators of condition-relevant niches. However, co-clustering methods have a limitation in that they cannot capture supervised information from control and condition slices. The STAGATE method utilizes a graph neural network to learn embeddings of cell niches. We configure the clustering parameters to align the final clusters with the known number of spatial domains with an additional placeholder cluster to place the condition-relevant niches. The CellCharter method leverages pre-trained single-cell latent vectors using scVI^63^ and spatial location data to delineate spatial domains. We maintain consistency with the specified clustering targets and adhere to the official tutorial’s steps to pretrain an scVI model and derive the final output. Since the CellCharter clustering number is automatically determined in its procedure, we limit the upper bound of the clustering number search procedure in CellCharter to match the STAGATE setting (original spatial domain number + additional placeholder class). For both STAGATE and CellCharter, since they are designed for spatial clustering to obtain niche labels across slices and are not designed for differential analysis and cannot output niches with condition specificity, to enable them to be comparable with Taichi, we regard the niche cluster labels that uniquely occur in the condition slice as the condition-specific niche.

The second category includes differential abundance (DA) analysis methods (such as MELD^17^ and HIDDEN^42^), traditionally used in single-cell data analysis to identify condition-relevant cell states. While these methods effectively utilize the supervision information from control and condition labels, they are based entirely on gene expressions and do not incorporate the spatial information inherent to spatial omics data. For the ‘HIDDEN’ method, the method autonomously generates the final condition-relevant cells. In contrast, ‘MELD’ produces a likelihood score as the initial model output.

The third category is supervised spatial domain identification for spatial transcriptomics, including the recently proposed CytoCommunity^43^. Such methods facilitate the identification of specific spatial domains between condition and control slices, integrating both spatial and conditional data to enhance detection accuracy. In our analysis, we employ the recently proposed CytoCommunity method to extract cell niches under the supervision of slice-level labels. We use the default parameters specified in the official codebase, designating the condition as label 1 and control as label 0. This labeling approach is utilized to train the CytoCommunity model. Additionally, the number of clusters is set in accordance with the spatial co-clustering setup, ensuring consistency with the spatial clustering comparison.

## Analysis of the mouse DKD Slide-seqV2 data

### Dataset overview

The study utilized seven replicates from 10 mm sagittal kidney sections obtained from both wild-type (WT) control and diabetic kidney disease (DKD) mice, with expert-annotated cell-type labels. We set the 26 DKD slices as the condition slices and the 34 WT slices as the control slices. The entire dataset was employed as input for the Taichi analysis. The output of Taichi in this dataset is to partition each real DKD slice (expected to be mixture of healthy and disease) into two virtual slices, i.e., slice containing DKD-relevant niche (DRN) and slice containing WT-consistent niche (WCN).

### Cell proportion comparison

To validate our hypothesis that the WCN niche in the Taichi outcome on the DKD slice is more similar to WT than the DRN niche, we conducted a comparative analysis of cell proportions and the Cell-Cell Interaction (CCI).

For the cell proportion comparison, we examine the similarity from two perspectives: first, the global cell proportion comparison, and second, the pairwise local comparison. The global cell proportion comparison entails calculating the cell frequency vector separately for all three areas (WT, WCN, DRN). Specifically, we summarize all the cell proportions (normalized into 0 to 1) in all WT, WCN, and DRN slices, and compare the WT cell proportion vector versus the WCN cell proportion vector, and the WT cell proportion vector versus the DRN cell proportion vector. For local similarity, we first calculate the cell proportion vector for each individual slice, obtaining separate groups of vectors for WT, WCN, and DRN. Next, we assess the inter-group pairwise similarity by comparing the WT group with the other two groups, as well as the intra-group pairwise similarity within the WT group itself. We calculate three types of pairwise similarity scores: Pearson correlation, Spearman correlation, and cosine similarity. Then we can obtain three distributions for each similarity measurement (i.e., WT intra-group similarity, WT vs. DRN inter-group similarity, and WT vs. WCN inter-group similarity). These scores help us evaluate the difference between the WT intra-group similarity versus the other two inter-groups based on distributional aspects, i.e., the WT vs. WCN inter-group similarity should be more similar to the intra-WT similarity groups compared with WT vs. DRN.

### CCI difference comparison

In the CCI difference comparison analysis, we use the *squidpy.gr.interaction_matrix* function to calculate the normalized interaction matrix for each group in both the DKD and WT slices. The interaction values within the matrix for any two cell types are derived from the total number of edges shared between those cell types in the spatial neighbors graph. This graph is constructed using the *squidpy.gr.spatial_neighbors* function with default parameters. For the DKD slices, the interaction matrix for the DRN and WCN groups is constructed based on a virtual slice defined by the Taichi results, while for the WT group, it is based on the original WT slices.

We analyze the interaction matrix from two perspectives. First, we assess the differences in interaction values by treating each interaction within the same group as one distribution, and then comparing these distributions using the two-sided rank-sum test. Given the datasets of 34 WT, 26 WCN, and 26 DRN interaction matrices, each element within these matrices represents a distribution with 34, 26, and 26 data points, respectively, reflecting the CCI conditions in these groups. We compare the WT distribution to the WCN and DRN distributions to determine if significant differences exist in the CCI comparisons between WT vs. WCN and WT vs. DRN. This process yields hypothesis test outcomes with p-values for each CCI.

Furthermore, since our goal is to evaluate the overall similarity in the CCI matrices— rather than considering each CCI individually—and because we need to control the False Discovery Rate (FDR) amid multiple CCI comparisons within the same matrix, we employ the Benjamini-Hochberg (BH) procedure. This adjustment method allows us to manage the FDR and to compare the outcomes of WT vs. WCN and WT vs. DRN based on the number of CCIs with significant differences (FDR adjusted p-value < 0.05) under controlled FDR conditions.

Moreover, we investigate the differences in CCI patterns between the DRN and WCN groups compared to the WT group. We employ the element-wise absolute difference between matrices as the measure of CCI variation. Specifically, for two CCI matrices, A and B, we calculate the matrix of absolute differences, *D* = |*A* − *B*|. Each element in D represents the difference for the corresponding CCI between the two matrices, with larger values indicating greater differences. We construct these difference matrices pairwise for WCN vs. WT and DRN vs. WT, resulting in 676 pairwise difference matrices for each group comparison. From these matrices, we generate distributions that delineate the difference patterns in each group for every CCI.

To determine which differences are statistically significant, we conduct a one-sided rank-sum hypothesis test on the difference distributions for WCN vs. WT and DRN vs. WT. We control the False Discovery Rate (FDR) using the Benjamini-Hochberg (BH) procedure. This analysis helps us identify which interactions in DRN vs. WT exhibit significantly larger differences (FDR adjusted p-values < 0.05) compared to WCN vs. WT.

### Spatial domain-specific genes and metagenes

After applying our Taichi procedure to segment the DRN/WCN on an original DKD slice, we selected a sample from the DKD slices to analyze the gene expression patterns between the DRN and WCN niches within the same slice. We aim to identify DRN-specific spatially variable genes (SVGs) and metagenes consistent with previous studies^64,65^.

We start by constructing the neighborhood set for each cell with a pre-defined radius circle around each cell in the DRN. Any cell from the WCN located within this circle is considered a neighbor. This radius is specifically chosen to ensure that, on average, each cell in the DRN has about 10 neighbors. Once defined, we collect all the neighbors of each cell in the DRN to create a neighboring set.

Following this, we conduct differential expression (DE) analysis between cells in the DRN and those in the WCN neighborhood set using the Wilcoxon rank-sum test. Genes that show a False Discovery Rate (FDR)-adjusted p-value of less than 0.05 are identified as candidates for SVGs.

To further refine our selection, we apply three additional criteria to ensure the genes have enriched expression patterns specifically in the DRN. First, the gene must be expressed in over 80% of the DRN cells, and the ratio of the percentages of cells expressing the gene in the DRN to those in the WCN must exceed 1. Then, the expression fold change for the gene between the DRN and WCN must be greater than 1.5. Only the genes in the previous SVG candidates that meet these three criteria can be regarded as final DRN-specific SVGs.

Subsequently, we computed the DRN-specific spatially variable metagenes. The objective is to identify a group of genes that, when combined, form a ‘meta gene’ exhibiting an enriched expression pattern in the DRN. To construct this meta gene, we employ a multi-step iteration procedure:

Initially, we relax the criteria for filtering SVGs by lowering the fold change threshold. From these SVGs, we randomly select two genes as the base genes denoted as *g*_0_ with the expression *X*_0_ in the entire slice. Subsequently, our goal is to enhance the spatial expression pattern in the DRN by aggregating expression from additional genes to these base genes.

To facilitate this, we first calculate the mean expression level of *g*_0_ in the cells of the DRN, denoted as *e*_0_. We then identify all cells from WCN that have an expression level exceeding *e*_0_ of *g*_0_, forming a control group. Using this setup, we conduct differential expression (DE) analysis comparing cells from the DRN against those in the control group using the Wilcoxon rank-sum test. The gene exhibiting the smallest FDR-adjusted p value with higher expression in the DRN is selected and denoted as *g*_0+_ with gene expression *X*_0+_. Similarly, we perform DE analysis on cells from the control group against those from the DRN. The gene that shows the smallest FDR-adjusted p value and is more highly expressed in the control group is identified as *g*_0-_ with gene expression *X*_0-_.

The expression level *X_mt_*_1_ of the meta gene *mt*_1_ is subsequently calculated based on the expression levels of these selected genes:

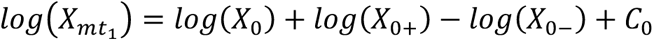

Where *C*_0_ is the constant to ensure the log value non-negative. Then, we can apply this procedure iteratively, and the metagene *mt_t_* in *t* iteration is the combination of the last *t* − 1 iteration *X_mt_*_(*t*-1)_ and the *X*_(*t*-1)-_, *X*_(*t*-1)+_ which are search by the same procedure of *g*_0_ but replace the base gene as *mt_t_*_-1_. Formally, it can be denoted as:

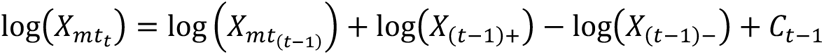

This iteration is expected to increase the difference of mean expression between the DRN and control groups selected in each iteration than last iteration and finally get the metagenes meet the criteria proposed in the previous single SVG selection. We here set the iteration limitation as 10 and stop the iteration when obtaining the metagene that meet the requirements.

### Spatial co-occurrence

The cluster co-occurrence ratio calculates a score reflecting how often specific clusters appear together in spatial dimensions. We use this metric to investigate a particular cell type of interest that conditions the co-occurrence calculation for all cell types. This calculation covers a series of radii, each with a specified diameter *d*, across the examined area. For this experiment, we employ the function in *squidpy.gr.co_occurrence* with default parameters. We specifically calculate the co-occurrence score conditioned on the kidney granular cell for all cell types in the given area. The outcome is a co-occurrence score for each cell type within a given diameter in space, where a higher score in a smaller diameter suggests closer physical proximity and interaction in the original area. Consequently, we independently calculated the co-occurrence scores for the WCN and DRN areas to examine the co-occurrence conditions based on the same kidney granular cell in these two distinct niches.

## Analysis of the Human CRC CODEX data

### Dataset overview

The dataset is characterized by extreme variations in intratumoral microenvironment (iTME) architecture within colorectal cancer (CRC). One group exhibited newly formed tertiary lymphoid structures (TLS) at the tumor’s invasive front, a phenomenon termed ‘Crohn’s-like reaction’ (CLR), whereas the other group lacked TLS, displaying diffuse inflammatory infiltration (DII) instead. Notably, patients with CLR demonstrated significantly longer survival times compared to those presenting with DII. From a database of 715 CRC patients, 17 patients with CLR and 18 with DII were selected for further analysis. This subgroup included 140 tissue regions from these 35 patients, each analyzed using 56 protein markers.

### Benchmarking by TLS detection

As outlined in the dataset overview, a key distinction between the DII and CLR subtypes is the presence of the TLS structure, annotated as ‘Follicle’ Cellular Neighborhood (CN) in the source data’s original publication. We have therefore set ‘Follicle CN’ as the positive label and are using recall as the metric to evaluate the model’s proficiency in detecting condition-specific niches within the DII samples. Specifically, we assessed the capability of both Taichi and baseline methods to identify DII-relevant niches, based on the golden label of TLS presence in the DII subtype. We focus solely on recall because, while other marker niches for the DII subtype may exist, a more effective method should at least accurately capture TLS in DII samples to demonstrate its capability in identifying relevant niches.

We implemented two baseline methods, CellCharter and CytoCommunity, recognized for their capability to manage large-scale datasets while preserving the spatial topological structure. For CellCharter, we adhered to the hyperparameters and pretraining method (trVAE) as recommended in its official guidelines. For CytoCommunity, we utilized the official default parameters, which include 10-fold training and the standard network parameters. For both methods, we set the clustering search upper bound/cluster number to match the original CN category number, which is 10. We evaluated the detection outcomes of these methods by focusing on areas that showed a significantly higher proportion (p<0.05) in CLR samples, demonstrating their detection efficacy. For Taichi, we used the expert-annotated cell types from the original source as input features. All methods were compared using the recall metrics previously mentioned to assess their performance in accurately detecting relevant niches.

## Analysis of the Head and Neck Cancer and Colorectal Cance clinical CODEX data

### Dataset overview

The Space-GM paper presents three CODEX slice datasets, each furnished with detailed, multi-label clinical annotations. The first dataset, UPMC-HNC, profiles head and neck cancer across 308 slices and includes annotations for evidence of disease, recurrence status, and HPV infection. The Stanford-CRC dataset comprises 292 colorectal cancer (CRC) samples, annotated with patient outcomes (alive/deceased), recurrence status, and a follow-up status of alive after 60 months or recurred/dead within 60 months. The DFCI-HNC dataset includes 58 slice samples, differentiated by low and high primary tumor responses to treatment (pTR)

### ROC curve

We utilized the fraction of Taichi’s condition-relevant niches area on each slice as a fine-grained indicator of intra-group heterogeneity within the original condition groups. This approach allowed for detailed examinations of these scores across specific condition groups. In the UPMC-HNC dataset, we apply Taichi using the evidence-of-disease as the condition label and employed the resulting area ratio as the score to compute the ROC-AUC for assessing both HPV infection and recurrence status. For the DFCI-HNC dataset, we designated a higher pCR (>10) as the condition label and used the area fraction scores to evaluate the correlation with continuous pCR scores across several thresholds (10-20, 20-30, 30-40, 40, 80-90, >90, 100). In the Stanford-CRC dataset, we utilize Taichi using the deceased condition label and calculated the ROC-AUC for the deceased/recurred within 60 months.

### Survival length curve

Additionally, we categorized the deceased group in Stanford-CRC into high-risk and low-risk subgroups based on the mean Taichi outcome condition-relevant niches area size and analyzed the significance of this stratification on survival length using Kaplan-Meier (KM) curves with log-rank test.

## Analysis of the Human Immunotherapy TNBC IMC data

### Dataset overview

The dataset employed Imaging Mass Cytometry (IMC) to examine the expression of 43 proteins at a subcellular level in tumor samples from patients with triple-negative breast cancer (TNBC) participating in the NeoTRIP randomized controlled trial. This trial evaluated the efficacy of neoadjuvant chemotherapy alone (carboplatin and nab-paclitaxel) versus chemotherapy combined with anti-PD-L1 immunotherapy (carboplatin, nab-paclitaxel, and atezolizumab) through 1:1 randomization. The final samples derived from 279 patients at three different stages: before treatment (baseline, n=243), and on the first day of the second treatment cycle (on-treatment, n=207). We analyzed all baseline state data, along with data from the chemical and immunotherapy co-process. Each cell was annotated by cell neighborhood (CN) and cell type (CT) in the original dataset, we employ the CT as the input feature for Taichi. Pathological complete response (pCR) data was used as the condition label, and Rd (Residual disease) as the control label.

### Tensor decomposition

Initially, we constructed frequency matrices for each cell type across different cell neighborhoods within Taichi-identified pCR-relevant and Rd-relevant niches. For both pCR-relevant and Rd-relevant niches, we summarized the frequency of each cell type in each cell neighborhood to form a CN-CT frequency matrix for each niche, which were then stacked to create a third-order tensor. Subsequently, we performed non-negative tensor decomposition using a Tucker Decomposition with a core tensor configuration of [2,2,5]. This decomposition process involves multiplying the core tensor by the cell type loader vector and the cell neighborhood loader matrix, resulting in a cell type and cell neighborhood interaction tensor. Mathematically, this process is described as follows:

Let *F_pCR_* and *F_Rd_* represent the CN-CT frequency matrices for pCR-relevant and Rd-relevant niches on entire dataset, respectively. These matrices are stacked to form the three-order tensor *T*. The tucker tensor decomposition is applied to *T* with the core tensor *C* defined as [2,2,5]. The decomposition results in a cell type loader matrix *L_ct_* and a cell neighborhood loader matrix *L_cn_*. The Taichi label Loader matrix *L_label_*. The interaction tensor *I* is then calculated by multiplying the core tensor *C* with *L_ct_* and *L_cn_*., which is expressed as:

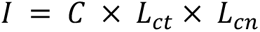

Each matrix within the tensor *I* represents the interaction between a cell neighborhood factor and a cell type factor, detailing their influence on each other separately. Then, we can use the *L_label_* matrix to assign the corresponding cell neighborhood factor-cell type factor interaction matrix for the pCR/Rd niches. Specifically, we utilize the column with the larger factor values in the *L_label_* matrix as the indicator for determining the belonging of the factor interaction matrix.

## Scalability Analysis

### Dataset overview

We simulate the large-scale dataset scenarios, S1 and S2, through duplicate simulations. For S1, we use the first layer of the previously studied MERFISH mouse brain slice, which contains 5,570 cells. We duplicate this MERFISH slice by factors of 2, 10, 100, and 1,000 to construct a multi-slice dataset. For the S2 scenario, we employ the MERSCOPE mouse brain dataset, specifically using replicate 1, which comprises 161,875 cells. We then duplicate this MERSCOPE slice by 2, 10, and 100 times to construct the S2 scenarios.

### Computational resources

All methods were run on the same machine, equipped with a 32-core Intel(R) Xeon(R) Gold 6338 CPU at 2.00 GHz and an Nvidia A100 (80G) GPU, to ensure fairness. We adhered to the default parameters for all methods, with the exception of adjusting the running epochs for scVI used in CellCharter to 100, to ensure consistency across different datasets. We focused our benchmarking on spatial information-aware methods.

### Computational resource

All experiments are performed on a server running Ubuntu with a 32-core Intel(R) Xeon(R) Gold 6338 CPU at 2.00 GHz and an Nvidia A100 (80G) GPU

### Data availability

Dataset 1: https://www.science.org/doi/full/10.1126/science.aat5691.

Dataset 2: https://datadryad.org/stash/dataset/ doi: 10.5061/dryad.8t8s248/.

Dataset 3-6: Generated from Dataset 2.

Dataset 7-9: https://vizgen.com/resources/.

Dataset 10: https://www.ncbi.nlm.nih.gov/geo/query/acc.cgi?acc=GSE190094.

Dataset 11: https://dx.doi.org/10.17632/mpjzbtfgfr.1.

Dataset 12-14: https://app.enablemedicine.com/portal/atlas-library/studies/92394a9f-6b48-4897-87de-999614952d94?sid=1168.

Dataset 15: https://doi.org/10.5281/zenodo.7990870.

### Code availability

The Python implementation and tutorial of Taichi is available at https://github.com/C0nc/TAICHI.

## Acknowledgments

Z.Y. acknowledges the support by Shanghai Municipal Science and Technology Major Project (No. 2018SHZDZX01), ZJ Lab, Shanghai Center for Brain Science and Brain-Inspired Technology, and 111 Project (No. B18015).

We acknowledge the research teams of “Spatial predictors of immunotherapy response in triple-negative breast cancer” for providing permission to use the large-scale immunotherapy TNBC dataset.

## Author contributions

Z.Y. and Y.C. conceived and designed the study, developed the computational methods, performed the analysis, and wrote the manuscript.

## Competing interests

The author declares no competing interests.

## Inclusion & Ethics

Not relevant.

---

**Supplementary Fig. 1 | Methods results in the MERFISH simulations**.

(a-h), Visualization of TP, TN, FP, FN in the spatial plots of different methods’ results on MERFISH simulations when the perturbation (at different rates) occurs in different niches, including medial preoptic area (MPA) (a), medial preoptic nucleus (MPN) (b), bed nuclei of the stria terminalis (BST) (c), columns of the fornix (fx) (d), paraventricular hypothalamic nucleus (PVH) (e), paraventricular nucleus of the thalamus (PVT) (f), third ventricle (V3) (g), and periventricular hypothalamic nucleus (PV) (h). The methods outputs were visualized in (i-p). Quantitative results of different perturbation rates were shown in (q).

**Supplementary Fig. 2 | Spatial clustering results**.

(a-c), Visualization of the spatial clustering results of CellCharter (a), CytoCommunity (b), and STAGATE (c) in both control and condition slices of STARmap simulations. The spatial clustering was performed in their multi-slice mode, to ensure labels from different slices are comparable. (d-f), Visualization of the spatial clustering results of CellCharter (d), CytoCommunity (e), and STAGATE (f) in both control and condition slices of MERFISH simulations. The spatial clustering was performed in their multi-slice mode, to ensure labels from different slices are comparable.

**Supplementary Fig. 3 | Visualization of Taichi results from the three tumor atlases.** Visualization of the Taichi-identified condition-relevant niches and associated cell types in three tumor atlases: (a) DFCI-HNC, (b) UPMC-HNC, and (c) Stanford-CRC.

## Notes

### Competing Interest Statement

The authors have declared no competing interest.

